# Determining susceptibility loci in triple negative breast cancer using a novel pre-clinical model

**DOI:** 10.1101/2024.02.08.579359

**Authors:** Samson Eugin Simon, Boston W. Simmons, Minjeong Kim, Sydney C. Joseph, Emily Korba, Sandesh J. Marathe, Margaret S Bohm, Sidharth S. Mahajan, Casey J. Bohl, Robert Read, Jeremiah R. Holt, D. Neil Hayes, Lu Lu, Robert W. Williams, Laura M. Sipe, David G. Ashbrook, Liza Makowski

**Author notes:** Co-first authors. Co-Corresponding authors: Liza Makowski, Cancer Research Building Room 322 UTHSC 19 S Manassas Street, Memphis, TN 38163, David Ashbrook, Coleman Building, Room E226B Department of Genetics, Genomics, and Informatics, UTHSC 956 Court Avenue, Memphis, TN 38103, Laura Sipe, Department of Biological Sciences, University of Mary Washington; 1301 College Avenue, Fredericksburg, Virginia 22401, USA. **Conflict of interest statement**: The authors have declared that no conflict of interest exists.

## Abstract

Breast cancer (BC) is the most common cancer and the second cause of death in US women. Our lack of understanding of how genetic variants affect molecular mechanisms that mediate BC aggression poses a substantial obstacle to advancements in cancer diagnosis and therapy. To examine genetic variants on BC traits, a novel murine model was created with robust phenotypic and genomic variation. The FVB C3(1)-T-antigen (“C3Tag”) mouse develops spontaneous tumors in the mammary glands of female mice with a mean latency of 4-5 months of age. This genetically engineered mouse model (GEMM) is well established to resemble human basal-like TNBC. TNBC is an aggressive subtype with few clinical approaches and poor patient outcomes. Thus, to model human heterogeneity in BC outcomes, we systematically crossed the C3Tag GEMM into the BXD recombinant inbred family – the largest and best characterized genetic reference population. The new model is termed “BXD-BC” and F1 hybrids of the cross have isogenic genomes that are reproducible. BXD-BCs are a potent tool to determine the impact of genetic modifiers on BC tumor traits. We hypothesized that examination of BXD-BC GEMMs will enable the identification of susceptibility loci, candidate genes, and molecular networks that underlie variation of multiple BC phenotypes. Using N=29 BXD-BC strains, we demonstrated significant heritable variations in the severity of TNBC characteristics such as tumor latency, multiplicity, and survival. Interestingly, 2 BXD-BC strains never developed tumors out to 1 year of age. Thus, BXD-BC strains demonstrate variance in cancer susceptibility and progression compared to the parent C3Tag GEMM, indicating the presence of genetic modifiers. Through an unbiased systematic quantification of breast cancer severity across BXD-BC hybrids, we identified several significant quantitative trait loci (QTL) and candidate genes for specific tumor traits. In combination with public human GWAS datasets, we defined syntenic regions, candidate genes, and underlying networks through cross-species systems genetics analyses to demonstrate the translational validity of conserved, biologically relevant, and targetable candidates. Our findings suggest conserved candidates predicting TNBC patient survival. In sum, the BXD-BC resource is an innovative, reliable, and robust preclinical model that reflects robust genetic heterogeneity. Using cutting edge systems genetics, we have identified genetic modifiers of BC phenotypic variation that could be targeted to advance therapeutic limitations or as biomarkers of risk or response to therapy.

## INTRODUCTION

Breast cancer (BC) is a significant global health concern, involving a heterogeneous group of malignancies that vary in their molecular characteristics, clinical behavior, and treatment responses. This is a grave clinical challenge for the ∼30,000 patients diagnosed with this disease annually in the United States. BC is the second most common cause of death in US women. Worldwide, BC ranks number one in cause of death in females ^1^. Our lack of understanding of how genetic variants affect molecular mechanisms that mediate BC aggression and impact effective anti-tumor therapies poses a substantial obstacle to advancement in cancer therapy. Triple-negative breast cancer (TNBC) represents around 15-20% of all BCs and is associated with aggressive clinical behavior, such as early metastasis, and limited treatment options when compared to other BC subtypes due to limited therapeutic options. Human studies have identified TNBC risk factors including environmental (e.g., age, obesity, alcohol) and genetic (e.g., BRCA1/2, among others) ^2–6^. While deleterious germline mutations in genes including *ATM, PTEN, HER2, BRCA1,* and *BRCA2* are commonly utilized in genetic diagnostic testing, they only explain approximately 10% of BC cases,^7^ emphasizing the complexity and limitations of diagnosis and risk prediction. Genetic influences on cancer involve various genetic modifiers impacting complex cellular and molecular processes. Together the interaction of modifier and causal genes govern the heterogeneity of cancer traits such as initiation, progression, metastasis, response to therapy, and risk of recurrence. Herein, we aimed to develop a novel murine model with robust phenotypic and heritable genomic variation in BC to identify gene variants that impact tumor aggression. Understanding the specific molecular mechanisms including patterns of gene expression and their underlying genetic modifiers that are driving TNBC tumor formation and aggressiveness will improve our understanding and treatment of the disease.

To create a novel pre-clinical BC model, we systematically crossed an established basal-like triple negative BC (TNBC) genetically modified mouse model (GEMM), the C3(1)-T antigen (“C3Tag”) model, with the best characterized murine genetic reference population—the BXD family – to create the novel “BXD-BC” resource. The C3Tag GEMM was created by Jeff Green at the NCI using recombinant expression of the simian virus 40 early region transforming sequences under the regulatory control of the rat prostatic steroid binding protein C3(1) gene in the FVB/N background strain ^8^. We and others have reported that the C3Tag GEMM is a reliably penetrant model: hemizygotes progressively develop intraepithelial neoplasia after ∼8 weeks, and ductal hyperplasia similar to ductal carcinoma in situ (DCIS) by ∼3 months of age ^9–13^, and finally adenocarcinoma by 5-6 months of age in 100% of the females ^9–14^. The C3Tag GEMM is a well-accepted model of the human basal-like TNBC subtype as it recapitulates the common loss of function mutations in the tumor suppressor genes retinoblastoma (Rb1) and TP53 ^15–19^. Thus, the C3Tag GEMM and human TNBC genetics align well ^8,14^. Taken together, the C3Tag GEMM is ideal for studying resultant changes in cancer onset and tumor parameters across genetic backgrounds.

Genetic modifiers of BC development may interact directly with cancer initiating genes (i.e. p53 and RB), influence cancer cell intrinsic mechanisms of DNA repair or cell cycle regulation, or alternatively impact response to cancer cells by the tumor microenvironment (TME) such as angiogenesis or antitumor immunity, to variably impact patient outcomes. The disadvantage of the isogeneic C3Tag mouse in the FV/N strain is the lack of genetic complexity that fails to represent typical human population variance, which limits interpretation and compromises the translational utility of discoveries ^20–22^. Therefore, we enhanced the C3Tag model by crossing it to the highly genetically variable BXD family. The BXD family is a large, genetically complex resource with matched multiscalar and multisystems phenome data ^20,23–25^. The BXD family is the largest rodent genetic reference population (GRP) and was generated over five decades ago by crossing C57BL/6J (B6 or “B”) and DBA/2J (D2 or “D”) – termed BXD – and subsequently inbreeding the F2 progeny for >20 generations ^26–30^. Each of the 152 fully inbred strains is genetically distinct and isogenic, i.e. fixed at each locus of the genome ^31^. The BXD, with two founder strains, has excellent power to detect loci with significant heritability using smaller cohorts compared to other recombinant inbred models such as the Diversity Outbred or the Collaborative Cross. To determine the impacts of genetic variants on TNBC, BXD females from multiple strains were crossed with C3Tag males resulting in BXD-BC F1 progeny in which females develop breast tumors. The novel BXD-BC GEMM combined mouse model now replicates the progressive tumorigenesis of human BC and the genetic variability of the human population. We present the BXD-BC GEMM as a highly valuable tool to discover insight that was previously intangible.

Herein, we report that BXD-BC F1 hybrids display greatly differing severity of presentation of TNBC phenotypes including traits such as tumor latency, multiplicity, and survival. Using unbiased systematic quantification of BC severity and heritability across BXD-BC hybrids, we identified heritable differences in distinct BC phenotypes that are driven by genetic modifier loci which segregate in the population. Through systems genetics analyses, we identified significant quantitative trait loci (QTLs) associated with BC phenotypes and expression-QTLs (e-QTLs) associated with tumor gene expression. Within the identified QTLs, candidate genes were narrowed following cross-species comparisons of existing human databases in syntenic regions. In sum, the BXD-BC hybrids demonstrate significant, heritable variation in tumor phenotypes which have yielded candidate genes using systems genetics. The generation of this reliable, reproducible, and robust BXD-BC resource will enable the discovery of genetic and environmental modifiers which will be leveraged to further understand BC.

## MATERIALS AND METHODS

### Reagents

All reagents were obtained from Sigma-Aldrich (St. Louis, MO) unless otherwise noted.

### Animals

Animal studies were performed with approval and in accordance with the guidelines of the Institutional Animal Care and Use Committee (IACUC) at the University of Tennessee Health Science Center (UTHSC, Animal Welfare Assurance Number A3325-01) and in accordance with the National Institutes of Health Guide for the Care and Use of Laboratory Animals. All animals were housed in a temperature-controlled facility with a 12-h light/dark cycle and *ad libitium* access to food and water. Mice were housed at UTHSC in the same animal facility to minimize external impacts. BXD nulliparous female breeders were supplied from the UTHSC colony maintained by Dr. Lu Lu and the Center for Integrative and Translational Genomics (CITG), directed by co-I Dr. Rob Williams. C3(1)-Tantigen (“C3Tag”) males were obtained by Jackson Labs, Inc. (Bar Harbor, ME) as strain #:013591 (*FVB-Tg(C3-1-TAg)cJeg/JegJ*; RRID:IMSR_JAX:013591) which are on the FVB/NJ genetic background. Mice were bred and maintained on Teklad Diets chow (Envigo #7912, Indianapolis, IN). Breeders were not maintained longer than 12 months of age. Sample size power calculations were carried out using the publicly available BXD power app, based on qtlDesign (Ashbrook DG 2019; Belknap 1998). C3Tag-BXD F1 hybrids were termed “BXD-BC” followed by the BXD strain ID #. The creation of BXD-BC model is represented in **Figure 1**.

**Figure 1.**
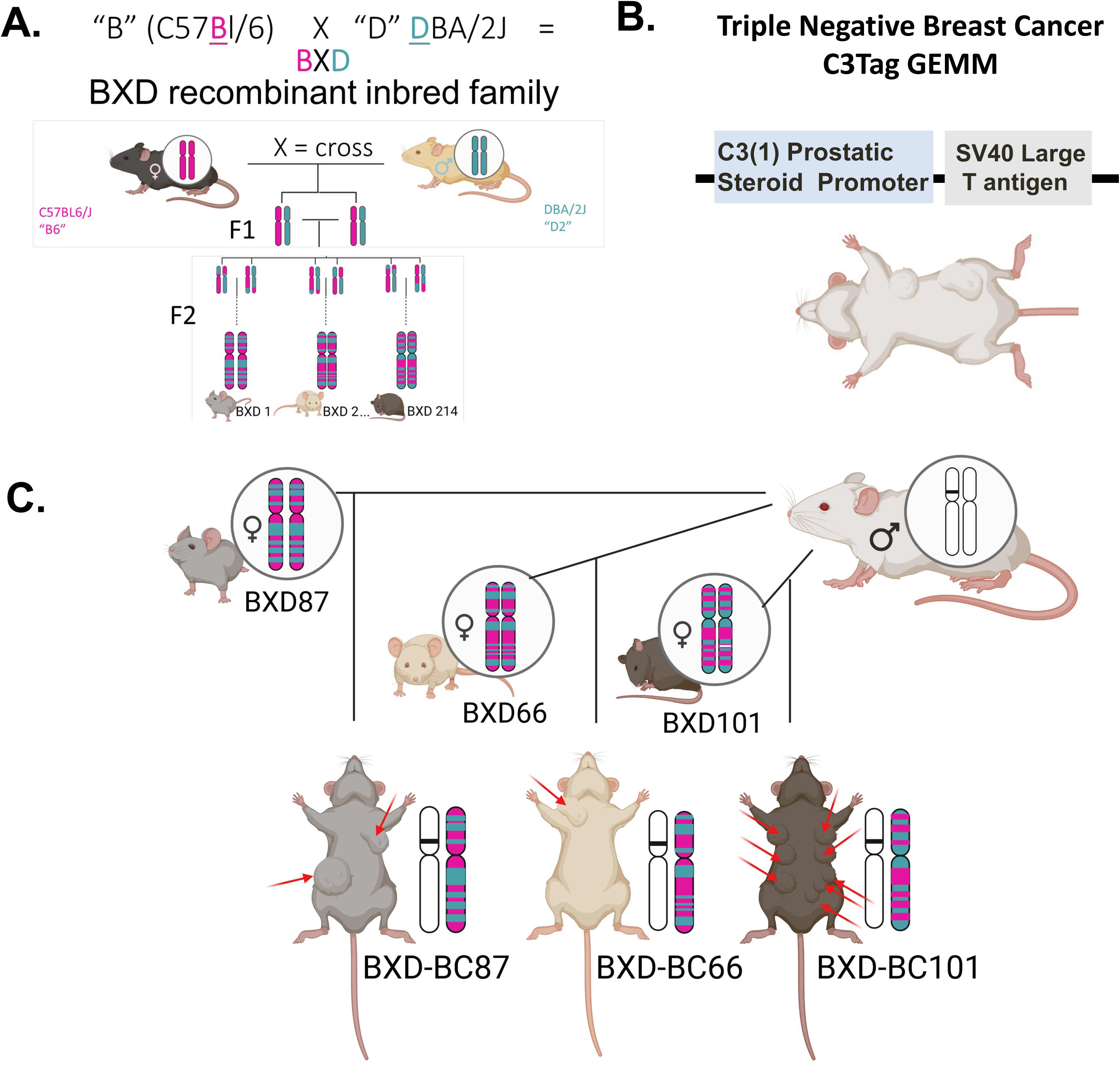
BXD-BC hybrids were generated by crossing C3Tag mice to various strains from BXD recombinant inbred family. **A**. C57BL/6J “B6” or “B” and DBA/2J “D2” or “D” parents were crossed to create BXD mice. F2 progeny of each BXD cross were iteratively inbred by sibling mating until generation F20+, at which point each BXD strain represents a unique mosaic of *B6* and *D2* alleles, homozygous at every loci. **B**. C3(1)-Tantigen (“C3Tag”) TNBC GEMMs were purchased from Jackson Labs and are well-established models of human basal-like TNBC breast cancer. The C3Tag are in the FVB/N background. **C**. Male mice transgenic for Tantigen were crossed into females of various strains of the BXD recombinant inbred family. C3Tag-BXD F1 hybrids were termed “BXD-BC” followed by the BXD strain ID #. Tumors develop spontaneously with 100% penetrance of transgene with latency, number, and size of tumor varying by mouse (red arrows in BXD-BC progeny).

### Tumor latency, progression, and endpoint

Latency for females C3Tag in the FVB/N genetic background has a mean of 19-20 weeks of age, with tumors arising as early as 11 weeks of age in rare cases depending on the variables tested ^9–13^, thus palpation for tumors was initiated at 9 weeks of age. Mice were palpated across all mammary glands twice weekly since tumors may develop in any mammary gland. Once palpable, tumors were measured by digital caliper three times weekly. The first tumor was called “T1” with subsequent tumors noted as “Tn+1.” Tumors were allowed to grow until a humane tumor endpoint was reached per IACUC protocol (1 tumor > 2cm in one direction or 3 tumors >1cm in diameter). Observable TNBC traits were recorded for over 13 phenotypes: age at latency; total tumor volume (mm^3^) at endpoint per mouse; volume, mass, or location of first tumor; total weight of all tumors dissectible (grams) at endpoint per mouse; age at endpoint; survival from latency to endpoint; location and number of tumors at sacrifice (multiplicity) per mouse.

### Endpoint

Mammary tumors were dissected from every mammary gland and weighed. In mice where no palpable tumors were present, the inguinal 4^th^ mammary gland was isolated. The T1 tumors or mammary glands were divided and flash frozen or formalin (10%), fixed, and paraffin embedded (FFPE) for histology. All additional tumors were flash frozen and stored at −80°C until analyzed.

### Histology and quantification

T1 tumors or unaffected 4th mammary fat pads were cut at 5 µm thickness. FFPE sections were stained with Hematoxylin and Eosin (H+E) and scanned by Thermo Fisher (Panoramic 250 Flash III, Thermo Fisher, Tewksbury, MA) scanner in the UTHSC Center for Cancer Research Imaging Shared Resource. Veterinary Pathologist Dr. Robert Read blindly scored H+E sections of tumor and surrounding mammary fat pad adipose tissue for hyperplasia, DCIS, and invasion, similar to our previous work(Qin et al. 2016).

### Heritability

Heritability is the proportion of phenotypic variance that is explained by genetic differences. Heritability was calculated for measured traits to determine the proportion of phenotypic variance explained by genetic effects, rather than environmental, technical or stochastic effects. Narrow-sense heritability (h^2^) was estimated as the fraction of variance explained by strains in a simple ANOVA model ^32^.

### GeneNetwork

GeneNetwork (GN) ID for the project is BXD_24401, with trait IDs noted in results. Strain mean values for all collected phenotypes were uploaded to GeneNetwork.org. Bioinformatics methods within GeneNetwork are established to identify candidates based on BC traits and our deep phenome of the BXD family ^3,23,33–36^.

### Quantitative trait loci mapping & definition of significance

Mean value per strain was calculated to reduce environmental variation, increasing the ability to detect causal quantitative trait loci (QTL). The BXD family has been produced in several ‘epochs’, using both standard F2 recombinant inbred methods, as well as advanced intercross recombinant inbred methods ^37^. This has led to both expected and unexpected kinship between BXD strains. This kinship between strains can lead to bias, as it breaks the expectations of previously used methods. The Linear mixed model-based methods allow correction for kinship and for other covariates. Therefore, two complementary methods were used in this study: QTL analysis using R software package R/qtl2 (Broman et al. 2019) and Genome-wide Efficient Mixed Model Association (GEMMA, ^3^, which is accessible in GeneNetwork.org). R/qtl2 ^38^ calculates empirical statistical significance using 5000 permutations of the data. A p < 0.05 was treated as significant, and a p < 0.67 was treated as suggestive. A p < 0.63 equates to one false positive per genome scan. However, the likelihood of any chromosome having a QTL on it is approximately 1 in 20 (i.e., p < 0.05) due to 20 chromosomes in mice. The likelihood of two independent traits sharing the same QTL location by chance is therefore much lower than p < 0.05. GEMMA was used to provide a −log(p) value between each marker and the phenotype. A −log(p) > 4 threshold is defined as significant. Both R/qtl2 and GEMMA approaches use a kinship matrix to mitigate relatedness between the strains to identify QTLs. A 1.5 –log(p) / LOD drop confidence interval is used as a reliable metric previously shown to contain a causative variant in BXD strains ^30,39^. Therefore, this interval was used to generate a list of candidate genes for each phenotype with a significant QTL.

### Candidate gene identification

Cis-eQTLs were next identified as a variant in the region associated with that gene’s expression to aid in determination of viable candidate genes. To determine expression for cis-eQTL, defined as a significant QTL within ∼5Mb of the gene, tissue relevant to tumor biology already in GeneNetwork including subcutaneous white adipose tissue, spleen, T-cells, leukocytes, neutrophils, and lymph node were queried. Next, BXD whole genome sequencing (WGS) data (Ashbrook et al. 2022) was utilized to identify potentially deleterious variants within the QTL region. All segregating variants within the QTL interval were submitted to the variant effect predictor (VEP) ^40^, with a focus on variants that impact protein expression or function, to aid in identification of viable candidate genes. Potentially deleterious variants or variants which impact protein function were identified using the “Consequence”, “IMPACT”, “SIFT” ^41,42^ and “BLOSUM62” ^43^ annotations.

### Human phenome wide association analysis study (PheWAS) translation

Human phenome wide associations (PheWAS) was conducted using summary statistics available online from existing studies ^44–50^. We used human PheWAS databases for all the candidate genes in our QTLs to detect genes with relevant human phenotype associations (e.g. cancer or immune phenotypes). The human genomic regions syntenic to the mouse QTL were assessed using GWASatlas (https://atlas.ctglab.nl/PheWAS) and PheWeb (http://pheweb.sph.umich.edu/) were accessed to determine if genes within a mouse QTL were associated with relevant phenotypes in humans.

### Statistics

For mouse and tumor traits, statistical differences between experimental groups were determined using Kaplan Meier tumor-free survival analysis, One-way or Two-way ANOVA, or Student’s *t*-test (as noted in figure legends) with Fisher’s LSD test for individual comparisons. Outliers were identified and excluded based on the ROUT method with Q=1%. For body weight, body composition, and tumor volume over time within animals, data was treated as repeated measures. All statistics were performed using statistical software within Graphpad Prism (Graphpad Software, Inc., La Jolla CA) or software packages noted above. All data are shown as Mean ± standard error of the mean (SEM) unless otherwise noted. P values less than 0.05 were considered statistically significant unless noted above for QTLs.

## RESULTS

### Generation of the BXD-BC Genetically Engineered Mouse Model

Females from the BXD recombinant inbred family (**Fig 1A**) and C3(1)-Tantigen “C3Tag” (**Fig 1B**) males were crossed to generate BXD-BC females used to study BC phenotypes (**Fig 1C**). Twenty-nine BXD-BC strains were generated with a minimum of N=2, maximum of N=14, and a mean of N=8 F1 isogenic hybrid progeny in each strain (**Table 1**). All traits examined are listed with GeneNetwork ID in **Supplemental Table 1**.

**Table 1.** BXD-BC Strains Generated. Strain ID and mice per strain are indicated.

| **BXD-BC Strain ID** | **# mice per strain** |
| --- | --- |
| 1 | 2 |
| 24 | 7 |
| 29 | 7 |
| 43 | 9 |
| 44 | 11 |
| 51 | 7 |
| 60 | 9 |
| 63 | 5 |
| 65 | 8 |
| 66 | 5 |
| 70 | 4 |
| 75 | 9 |
| 78 | 10 |
| 79 | 7 |
| 83 | 9 |
| 87 | 10 |
| 89 | 9 |
| 90 | 7 |
| 101 | 5 |
| 102 | 12 |
| 113 | 14 |
| 124 | 12 |
| 154 | 4 |
| 161 | 11 |
| 210 | 5 |
| 214 | 11 |
| 128a | 9 |
| 73b | 8 |
| FVB | 6 |

### BXD-BC Tumor Phenotypes Show Phenotypic Heterogeneity Relative to C3Tag GEMM

The range for tumor latency for the FVB/N C3Tag GEMM is ∼19 weeks of age, but some tumors occur as early as 11 weeks of age. Therefore BXD-BC females were palpated starting at 9 weeks of age. Mice were palpated across all mammary glands two times weekly. Once palpable, tumors were measured by digital caliper three times weekly until the IACUC defined endpoint was reached. Multiplicity represented as the total number of tumors per mouse was quantified at endpoint and showed significant variance across BXD-BC strains (**Fig 2A**). The parental C3Tag strain (“FVB”) displayed a relatively low mean of 3.17 ± 0.91 tumors per mouse with a range from N=1 to 7 per mouse (**Fig 1A**). Strain BXD-BC210 developed the fewest tumors per mouse with a mean of 2.4 ± 0.68 tumors compared to the BXD-BC63 strain which had 4-fold greater tumor multiplicity compared to the BXD-BC210 strain, with a mean of 8.8 ± 1.66 tumors per mouse (**Fig 1A**). The average coefficient of variation (CV) across strains was 40.1%, but some strains had as low as 10.9% for BXD-BC1 while C3Tag parental FVB strain had the highest CV of 70.4% for multiplicity. Importantly, two strains never developed BC tumors, strains BXD-BC79 and BXD-BC51. Mice in these strains were maintained for 12 months, well past typical C3Tag mean latency in the FVB strain of ∼4-5 months of age. No mammary tumors developed in mice of these strains as confirmed by palpation thrice weekly and with dissection at endpoint. Of note, these mice did develop other rare tumors reported in the C3Tag strain such as mixed eccrine sweat gland tumors of the paw ^8^, confirming the cross occurred correctly in addition to accurate genotyping. In all strains that developed mammary tumors, the total tumor burden (or total mass) was quantified by weighing all identified tumors at endpoint (**Fig 2B**). Burden revealed a 3-fold variance across BXD-BC strains ranging from about 1.13 grams per mouse in BXD-BC66 to a mean of 3.25 grams per mouse in BXD-BC101. The C3Tag FVB parental strain displayed a mean of 3.01 with a range from 2.26 to 3.95 grams per mouse. The average coefficient of variation (CV) across strains was 42.0%, but some strains displayed a CV as low as 15.26% for BXD-BC1 while BXD-BC90 had the highest CV of 71.5% for total tumor burden.

**Figure 2.**
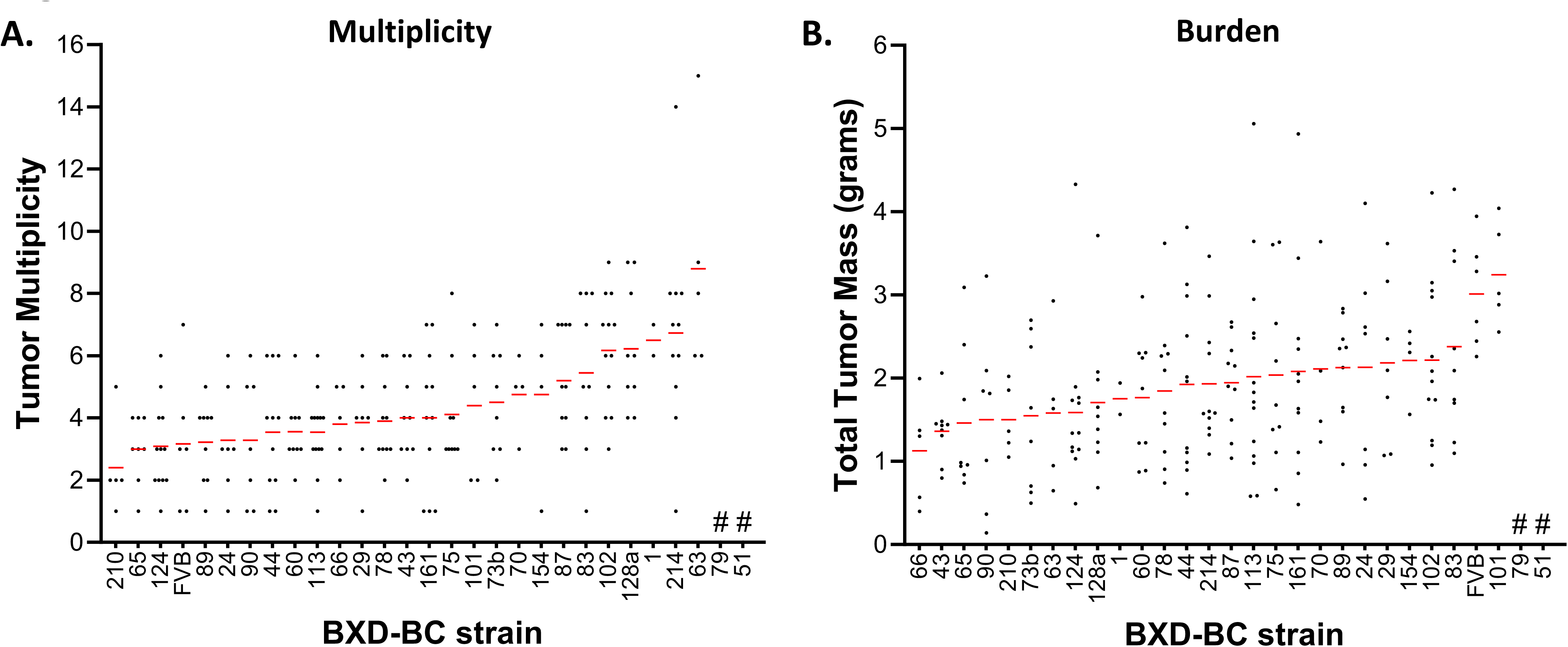
Novel BXD-BC hybrids demonstrate variability in tumor burden. Tumor phenotypes are ranked by mean (red line) with individual mice represented as dots. BXD strain number is on X axis for each BXD-TNBC hybrid. Parent C3Tag FVB/N strain is indicated as “FVB”. **A**. Tumor multiplicity (number of tumors/mouse). **B**. Tumor burden of all tumors (g/mouse). Note: # denotes strains with no tumor development after 12 months.

To examine tumor progression, age at latency (time of first tumor palpated) was recorded along with age at endpoint. **Figure 3** shows both latency and endpoint to represent progression reaching IACUC endpoint. Latency and endpoint vary by about 1.60 – 1.75-fold from the earliest onset strains (early latency) to the latest onset strains (delayed latency). The strains with the earliest latency included BXD-BC161, 66, 29, and 70 with rapid tumor onset before 18 weeks of age, suggesting an aggressive tumor phenotype. Strains that displayed very delayed latency included BXD-BC44, 210, 113, and 24 with tumor onset occurring past 23 weeks of age. Absolute progression was on average 4.77 weeks, with some strains rapidly progressing to endpoint such as BXD-BC90 at 3.06 weeks until endpoint, compared to others which had a much slower progression including BXD-BC154 and BXD-BC24 with progression times as long as 7.79 and 8.39 weeks, respectively. Overall, the C3Tag GEMM parental strain fell within the middle for both latency and endpoint, demonstrating that crossing into various BXD strains introduced genetic variants that either drove aggression or reduced aggression.

**Figure 3.**
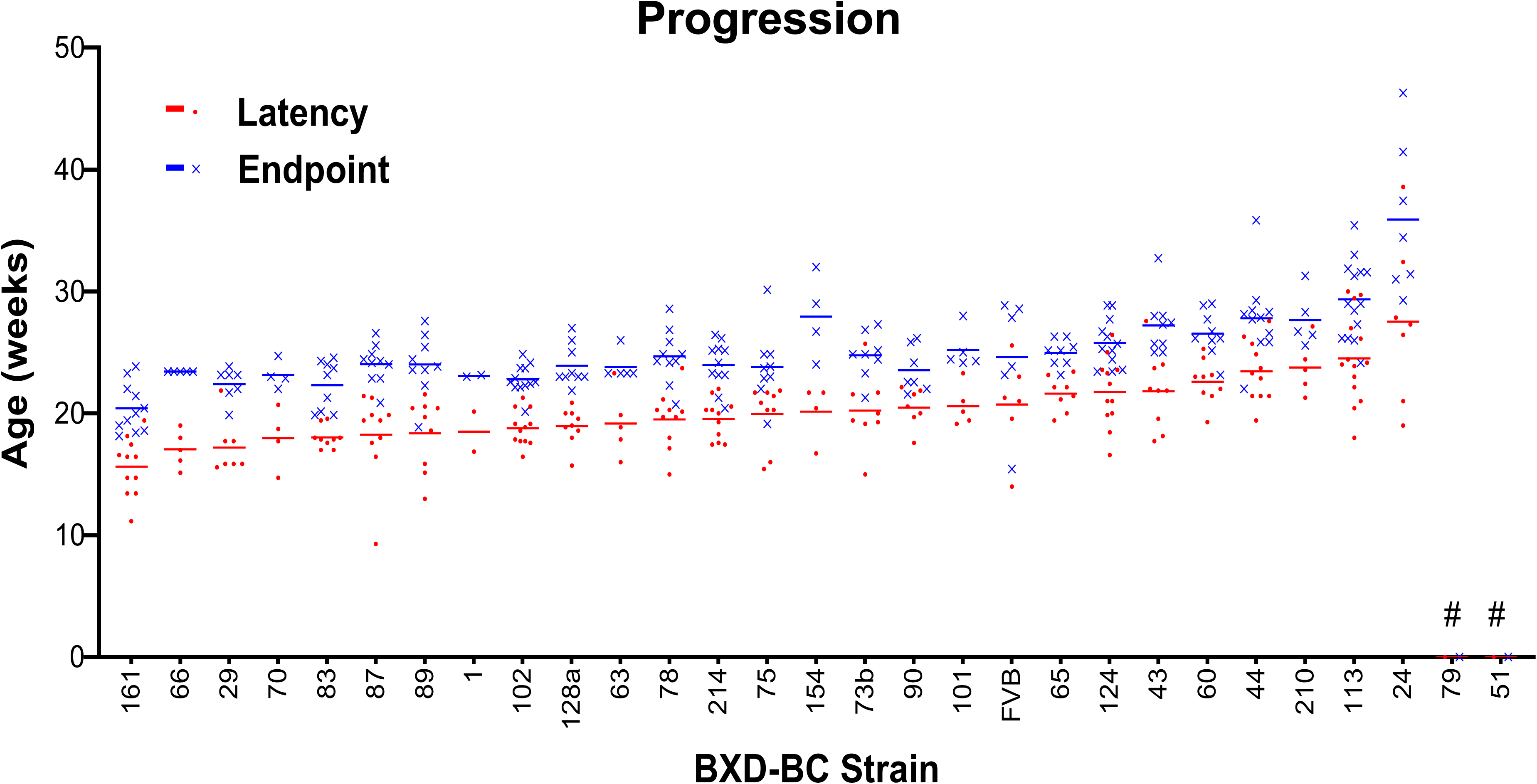
Tumor latency and endpoint are modified compared to parent strain in BXD-BC hybrids. Tumor phenotypes are ranked by mean (red line) for latency. BXD strain number is on X axis for each BXD-TNBC hybrid. Parent C3Tag FVB/N strain is indicated as “FVB”. Tumor latency is shown as black circle representing each mouse and the mean for that strain in red line while endpoint is indicated by grey “x” and blue line for the mean. Tumor progression is defined from weeks old at latency to endpoint. Note: # denotes strains with no tumor development after 12 months.

### Multiple Tumor Traits Display Significant Heritability in BXD-BC Strains

Heritability (h^2^) and significance of strain effect were calculated. The BXD-BC model demonstrated large, heritable variance in multiple tumor traits. In descending order, age at endpoint (h^2^=0.63), age at latency (h^2^ = 0.53), tumor multiplicity (h^2^=0.36), and survival (endpoint minus latency, h^2^=0.20) revealed robust and significant heritability, with p-values in **Table 2**. Importantly, traits describing the first tumor (T1) including the T1 mammary gland location, T1 tumor weight, and T1 tumor volume did not display significant heritability. Heritability calculations including the two BXD-BC strains that never developed tumor, BXD-BC51 and BXD-BC79, are shown in **Supplemental Table 2** which revealed that survival and age at endpoint are extremely heritable (h2=0.97 and 0.93, respectively) when these strains were included. Other traits display similar or relatively similar effects as those when the two strains are not included.

**Table 2.** Multiple Tumor Traits Display Significant Heritability in BXD-BC Strains. Heritability (h2) and significance of strain effect (p) are shown tumor traits collected for N=26 BXD-BC strains, with an average of 8 replicates per strain. Strain effect was tested by ANOVA. Strains that did not develop tumors were excluded. Bold indicates significance P<0.05.

| Phenotype (Trait) | $h^2$ | p-value |
| --- | --- | --- |
| Age at endpoint | 0.633 | **2.94E-28** |
| Tumor latency | 0.524 | **9.63E-19** |
| Multiplicity | 0.357 | **1.54E-08** |
| Survival (Time from latency to endpoint) | 0.194 | **0.022** |
| Endpoint 1 tumor >2cm | 0.152 | 0.172 |
| Endpoint 3 tumors >1cm | 0.152 | 0.174 |
| Total tumor weight | 0.145 | 0.228 |
| T1 tumor location | 0.138 | 0.359 |
| T1 tumor weight | 0.100 | 0.755 |
| T1 volume | 0.077 | 0.940 |
| Total tumor volume | 0.074 | 0.955 |

### Histologic Analysis of Tumors Revealed Variability in Cancer Traits, but Heritability Was Generally Low

Tumors were stained by H+E and analyzed by a veterinary pathologist, Dr. Robert Read, DVM. Analysis for each trait was averaged from over 3 randomly selected regions of interest in tumor H+E sections and a score was generated. Mitosis per high powered field were quantified as markers of aggressive tumors (**Fig. 4A**). Mitoses varied from 10-80 mitotic nuclei per high powered field. Epithelial to mesenchymal transition (EMT) type histology was scored from 0 to 4 with areas of no pleomorphism to faint to distinct sarcomatous transition. EMT was variable across strains with means ranging from 1-2.7 in each strain (**Fig. 4B**). Vascularity and stromal content were scored from 0-5 with areas of no presence to mild to scirrhous presence. This trait was the moderately variable across strains although most tumors displayed a score of 2 on average (**Fig. 4C**). Likewise, necrosis of tumor was scored from none to cavitary tumor loss across a five-point scale. This score was also moderately variable across strains with a mean of 2.4 (**Fig. 4D**). While some of the traits displayed variability across strains, the highest heritability was for mitosis per hpf and vascularity and stroma with an h2 of 0.34 and 0.33, respectively, with no significant strain effect for any trait (**Supplemental Table 3**). However, QTL for necrosis and vascularity/stroma content phenotypes are regularly mapped in the BXD population.

**Figure 4.**
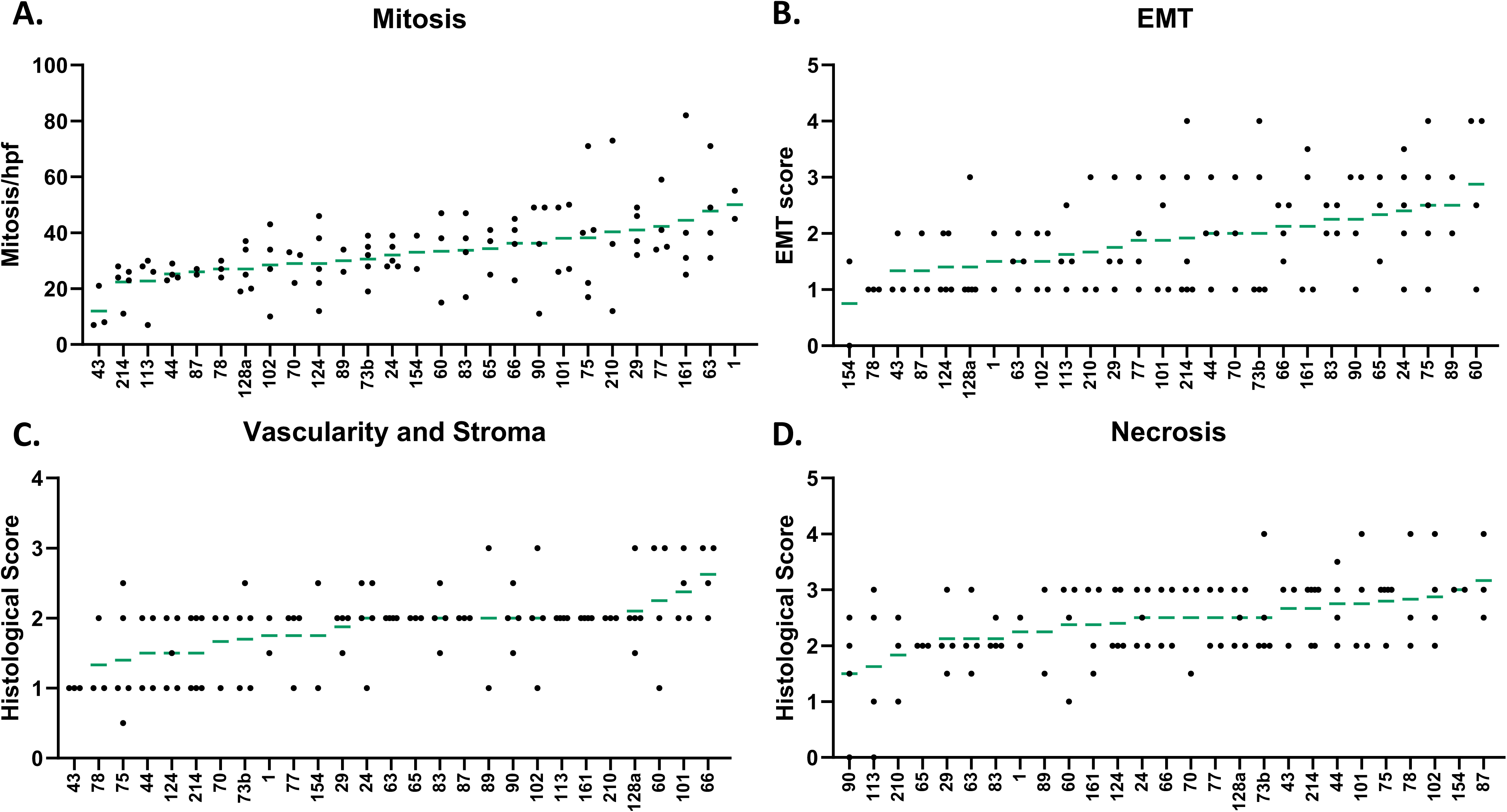
Tumor histologic analysis revealed variability phenotypes across strains. Tumors were stained by H+E and analyzed by a veterinary pathologist, Dr. Robert Read, DVM. Analysis for each trait was averaged from over 3 randomly selected regions of interest in tumor H+E sections and a score was generated. Data are ranked by mean (green line) with each dot representing 1 mouse tumor. **A**. Mitosis per high powered field (hpf) were quantified in 40X images. **B**. Epithelial to mesenchymal transition (EMT) type histology was scored from 0 to 4: 0=no pleomorphism; 1=subepithelial pleomorphism; 2=faint streaming; 3=distinct frequent streaming; and 4=distinct sarcomatous transition. **C**. Vascularity and stromal content were scored from 0-5: 0=none; 1=minimal stroma; 2=mild; 3=moderate; 4=heavy; and 5=scirrhous. **D**. Necrosis of tumor was scored from none to cavitary tumor loss across a five point scale.

### A QTL on Chromosome 16 was Identified for Tumor Multiplicity

To identify a critical region of the genome relative to variation in a trait, QTL mapping was undertaken using two complementary methods including GEMMA in GeneNetwork ^24,51,52^ and R/qtl2 in the R environment ^38^. For tumor multiplicity (GN ID BXD_ 24403), different QTL loci were identified dependent upon if all strains were used (i.e. including two strains that never developed tumors and treating them as zero), or without these two strains (treating them as N/A) (**Fig 5A-C**). Using all strains, 4 QTLs above the suggestive threshold (p < 0.67) were identified on Chr5 (peak 92.81 Mb), Chr10 (peak 120.59 Mb), Chr12 (peak 116.78 Mb), and Chr18 (peak 68.19 Mb), with the QTL on chr12 having the highest LOD value (3.12, p = 0.0678) (**Fig 5A**). When mapped removing the strains that never developed tumors, four suggestive QTLs were identified on Chr5 (peak 86.59 Mb), Chr8 (peak 10.77 Mb), Chr16 (peak 36.61 Mb) and Chr18 (peak 68.19 Mb), with the highest LOD being on chr16 (LOD = 3.24, p = 0.12) (**Fig 5B**). Because several loci appear to contribute to the phenotype, reducing the power to detect any single loci, remapping occurred controlling iteratively for the peak marker in each of these suggestive QTLs. When all samples were included, no loci reached the significance threshold. However, when the strains that never developed tumors were excluded and analysis included only tumor bearing mice, controlling for the chr5 QTL, the chr16 QTL became significant (LOD= 4.034, p = 0.0042) (**Fig 5C**). When the chr16 genotype was controlled for, the chr5 QTL became stronger (p = 0.056, LOD changed from 2.56 to 3.152, data not shown). **Figure 5D** demonstrates that the three genotype positions with the highest LOD scores (on chr5, chr12, chr16) the strains that develop the most tumors (n < 5.4) have the “B” or C57BL/6 genotype at all three loci, whereas the two strains that never develop tumors have the “D” or DBA2/J genotype at the chr5 and chr12 QTLs, and the B genotype at the chr16 QTL. These data suggest that several loci contribute to multiplicity in the BXD-BC model, yet the additive or interactive nature of loci remains unclear.

**Figure 5.**
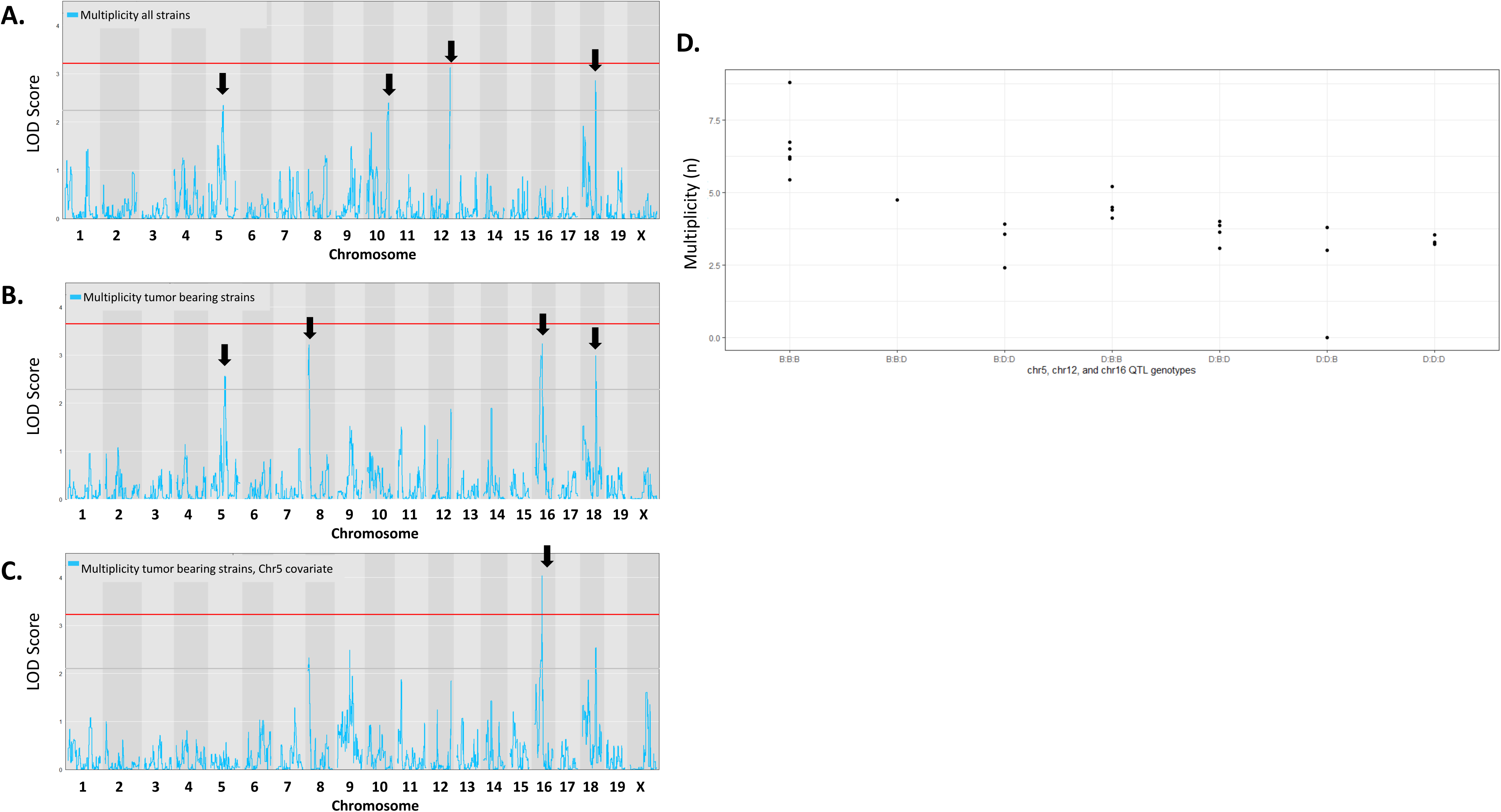
Genome wide QTL mapping for tumor multiplicity indicated significant QTL on chromosome 16. QTLs associated with tumor multiplicity were examined in R/qtl2. Genome wide QTL plot demonstrates the logarithm of the odds (LOD score (blue line) at each marker across the genome with chromosomes noted on the X – axis. The genome-wide significant threshold is indicated by red line (p < 0.05 calculated from 5,000 permutations of the data). The genome-wide suggestive threshold of p = 0.63 is indicated by grey line. **A.** Multiplicity in all strains was used to identify QTLs on Chr5 (peak At 92.81 Mb), Chr10 (120.59 Mb), Chr12 (116.78 Mb) and Chr18 (68.19 Mb), with the QTL on chr12 having the highest LOD value (3.12, p = 0.0678). **B.** Multiplicity in only tumor bearing strains was used to identify QTLs on Chr5 (86.59 Mb), Chr8 (10.77 Mb), Chr16 (36.61 Mb) and Chr18 (68.19 Mb), with the highest LOD being on chr16 (LOD = 3.24, p = 0.12). **C**. Chr5 peak marker was included as a covariate revealed a significant QTL present on chr16 (p = 0.0042, LOD 4.034). **D**. Tumor multiplicity (n) was plotted against the genotype at the peak markers on chr5 (92.81 Mb), chr12 (116.78 Mb) and chr16 (36.61 Mb). The three chromosome genotypes are presented in order, with B representing being homozygous with the B6-like C57Bl/6 allele, and D representing being homozygous for the D2-like DBA2/J allele on the X-axis.

### Gene Variants and Expression QTL (eQTL) Analysis Identified Candidate Genes associated with Chromosome 16 and Tumor Multiplicity

Gene variants can modulate phenotypes by two main routes – by changing the function of the gene product (e.g., a non-synonymous variant) or by altering the expression of the gene product. To investigate genetic variants, we used the Genome MUSter Web service ^53^. Using the chr16 QTL for tumor multiplicity, the 1.5 LOD drop confidence intervals were chr16: 33.75-36.91 Mb. To maximize the potential to detect a biologically significant gene in the QTL identified, the candidate genes were narrowed down to only protein coding. The chr16 QTL interval associated with tumor multiplicity contained 80 annotated genes and open reading frames, 42 of which were protein coding (**Table 3**). Of the protein coding candidate genes in the Chr16 QTL, there were 31 gene variants predicted to alter protein function or splice regions in 14 genes (*Ccdc14*, *Cd86*, *Dirc2*, *Dtx3l*, *Gm49662*, *Hspbap1*, *Mylk*, *Parp14*, *Parp9*, *Ptplb*, *Sema5b*, *Slc15a2*, *Slc49a4*, and *Wdr5b*).

**Table 3.** Chromosome 16 QTL Protein Coding Genes associated with Tumor Multiplicity. An interval generated in GEMMA for chr16 (33.75-36.91 Mb) contains 80 annotated genes and open reading frames (ORFs), of which 42 were protein coding with gene symbol, description, start point, length, and variant or single nucleotide polymorphims (SNP) counts and density reported.

| Symbol | Gene Description | Mb Start | Length (Kb) | SNP Count | SNP Density |
| --- | --- | --- | --- | --- | --- |
| *Heg1* | heart development protein with EGF-like domains 1 | 33.684384 | 87.19 | 11 | 0.13 |
| *Muc13* | mucin 13, epithelial transmembrane | 33.794037 | 25.89 | 1 | 0.04 |
| *Itgb5* | integrin beta 5 | 33.829665 | 119.67 | 152 | 1.27 |
| *Umps* | uridine monophosphate synthetase | 33.954782 | 12.26 | 4 | 0.33 |
| *Kalrn* | kalirin, RhoGEF kinase | 33.969073 | 604.20 | 1186 | 1.96 |
| *LOC110120412* | VISTA enhancer mm268 | 34.04964 | 1.04 | 2 | 1.92 |
| *Ropn1* | ropporin, rhophilin associated protein 1 | 34.649832 | 28.78 | 70 | 2.43 |
| *Ccdc14* | coiled-coil domain containing 14 | 34.690557 | 34.65 | 11 | 0.32 |
| *Mylk* | myosin, light polypeptide kinase | 34.745199 | 257.24 | 402 | 1.56 |
| *Hacd2* | 3-hydroxyacyl-CoA dehydratase 2 | 35.022421 | 86.77 | 103 | 1.19 |
| *Adcy5* | adenylate cyclase 5 | 35.154494 | 151.25 | 19 | 0.13 |
| *Sec22a* | SEC22 homolog A, vesicle trafficking protein | 35.311131 | 52.80 | 7 | 0.13 |
| *Pdia5* | protein disulfide isomerase associated 5 | 35.397308 | 93.61 | 110 | 1.18 |
| *Sema5b* | sema domain, seven thrombospondin repeats (type 1 and type 1-like), tr.. | 35.541154 | 123.11 | 211 | 1.71 |
| *LOC110120554* | VISTA enhancer mm1471 | 35.584523 | 5.25 | 15 | 2.86 |
| *Slc49a4* | solute carrier family 49 member 4 | 35.694062 | 75.31 | 175 | 2.32 |
| *Hspbap1* | Hspb associated protein 1 | 35.769726 | 58.74 | 10 | 0.17 |
| *Parp14* | poly (ADP-ribose) polymerase family, member 14 | 35.831892 | 39.63 | 56 | 1.41 |
| *Dtx3l* | deltex 3-like, E3 ubiquitin ligase | 35.926515 | 12.51 | 55 | 4.40 |
| *Parp9* | poly (ADP-ribose) polymerase family, member 9 | 35.93847 | 34.15 | 202 | 5.91 |
| *Kpna1* | karyopherin (importin) alpha 1 | 35.983288 | 53.85 | 14 | 0.26 |
| *Wdr5b* | WD repeat domain 5B | 36.04119 | 1.78 | 0 | 0.00 |
| *Fam162a* | family with sequence similarity 162, member A | 36.043844 | 27.72 | 3 | 0.11 |
| *Ccdc58* | coiled-coil domain containing 58 | 36.07166 | 20.46 | 13 | 0.64 |
| *Csta1* | cystatin A1 | 36.119946 | 11.24 | 7 | 0.62 |
| *Stfa2l1* | stefin A2 like 1 | 36.156811 | 5.14 | 19 | 3.70 |
| *Cstdc4* | cystatin domain containing 4 | 36.184212 | 3.90 | 0 | 0.00 |
| *Csta3* | cystatin A family member 3 | 36.210403 | 7.39 | 0 | 0.00 |
| *Csta2* | cystatin A family member 2 | 36.221562 | 35.89 | 4 | 0.11 |
| *Stfa1* | stefin A1 | 36.277148 | 8.22 | 10 | 1.22 |
| *Cstdc3* | cystatin domain containing 3 | 36.305357 | 7.36 | 0 | 0.00 |
| *Cstdc6* | cystatin domain containing 6 | 36.321665 | 12.67 | 9 | 0.71 |
| *Cstdc5* | cystatin domain containing 5 | 36.359382 | 8.19 | 1 | 0.12 |
| *Stfa2* | stefin A2 | 36.403946 | 4.42 | 3 | 0.68 |
| *Stfa3* | stefin A3 | 36.450537 | 4.86 | 26 | 5.36 |
| *Casr* | calcium-sensing receptor | 36.490585 | 71.56 | 77 | 1.08 |
| *Cd86* | CD86 antigen | 36.568956 | 97.20 | 4 | 0.04 |
| *Ildr1* | immunoglobulin-like domain containing receptor 1 | 36.693978 | 32.83 | 1 | 0.03 |
| *Slc15a2* | solute carrier family 15 (H+/peptide transporter), member 2 | 36.750161 | 35.00 | 1 | 0.03 |
| *Eaf2* | ELL associated factor 2 | 36.792884 | 82.18 | 7 | 0.09 |
| *Iqcb1* | IQ calmodulin-binding motif containing 1 | 36.82836 | 44.36 | 5 | 0.11 |
| *Golgb1* | golgi autoantigen, golgin subfamily b, macrogolgin 1 | 36.875093 | 57.99 | 9 | 0.16 |

Expression of genes within the ch16 QTL associated with tumor multiplicity were next identified through cis-eQTL analysis. To investigate variants which altered gene expression, we used existing transcriptome BXD family datasets in GeneNetwork to determine which genes within the QTL interval also had a *cis*-eQTL. *Cis*-eQTLs were analyzed in a number of tissues relevant to tumor or immune biology: T-helper cells, regulatory T cells, leukocytes, neutrophils, spleen, and white adipose tissue (because the mammary gland contains abundant adipose tissue). Cis-eQTL analysis revealed 8 genes with *cis*-eQTLs mapping to the chr16 QTL region: *Dirc2*, *Dtx3l*, *Stfa3*, *Zdhhc23*, *Kiaa0226*, *Hspbap1*, *Popdc2*, and *Slc15a2*. **Supplemental Table 4** summarizes these findings with the –logP score for latency, and Pearson’s r and Spearman’s rho correlations of gene expression and significance in indicated tissue types with tumor multiplicity are reported. The highest –logP scores were for *Slc15a2* and *Kiaa0226* in T helper cells and spleen (-logP score 10.2 and 10.1, respectively) and neutrophils with a –logP score 8.4 for *Slc15a2* and 9.6 for *Kiaa0226*, suggesting a role for these genes regulating immune cells in modulating tumor multiplicity.

### Gene Variants and Expression QTL (eQTL) Analysis Identified Candidate Genes associated with Chromosome 10 QTL and Tumor Latency

Including the two strains that did not develop tumors BXD-BC51 and BXD-BC79, a QTL for tumor latency (Genenetwork (GN) ID BXD_24401) was identified with –logP = 4.46 on chromosome 10 (Chr10) with a logarithm of odds (LOD) score of 4.53 (**Fig 6A**). The 1.5 LOD drop confidence interval covers chr10:117.295 – 121.534 Mb in GEMMA and 120.587 – 121.438 Mb in R/qtl2. The QTL on chr10 associated with tumor latency contained 12,466 gene variants in the BXD family in the larger GEMMA QTL confidence interval, and 1,866 in the smaller R/qtl2 confidence interval. The longer interval generated in GEMMA contained 121 annotated genes and open reading frames (ORFs), of which 27 were protein coding (**Table 4**). The latter smaller interval defined by R/qtl2 contained 13 genes, of which 6 are protein coding and include *Msrb3*, *Lemd3*, *Wif1*, *Tbc1d30*, *Gns*, and *Rassf3* (which are also in the overlapping GEMMA identified interval). Of note, two predicted missense variants were identified in *Tbc1d30* and a splice region variant in *Gns*. Expression of genes within the ch10 QTL associated with tumor latency was identified through cis-eQTL analysis. Examination of *cis*-eQTLs across tissues relevant to tumor or immune biology as analyzed above, including T-helper cells, regulatory T cells, spleen, and white adipose tissue. Of the 6 protein coding candidate genes in the ch10 QTL, three genes were identified with a significant *cis*-eQTL in at least one of these tissues: *Cand1*, *Tbc1d30*, and *Rassf3*. **Supplemental Table 5** summarizes these findings with the –logP score for multiplicity, and correlations of gene expression in indicated tissue types with correlations for latency and multiplicity. The highest –logP scores were for *Tbc1d30* and *Rassf3* in spleen (-logP score 10.5 and 7.9, respectively) and T regulatory cells (-logP score 7.9 and 4.3, respectively) suggesting a role for these genes regulating immune cells in modulating tumor latency.

**Figure 6.**
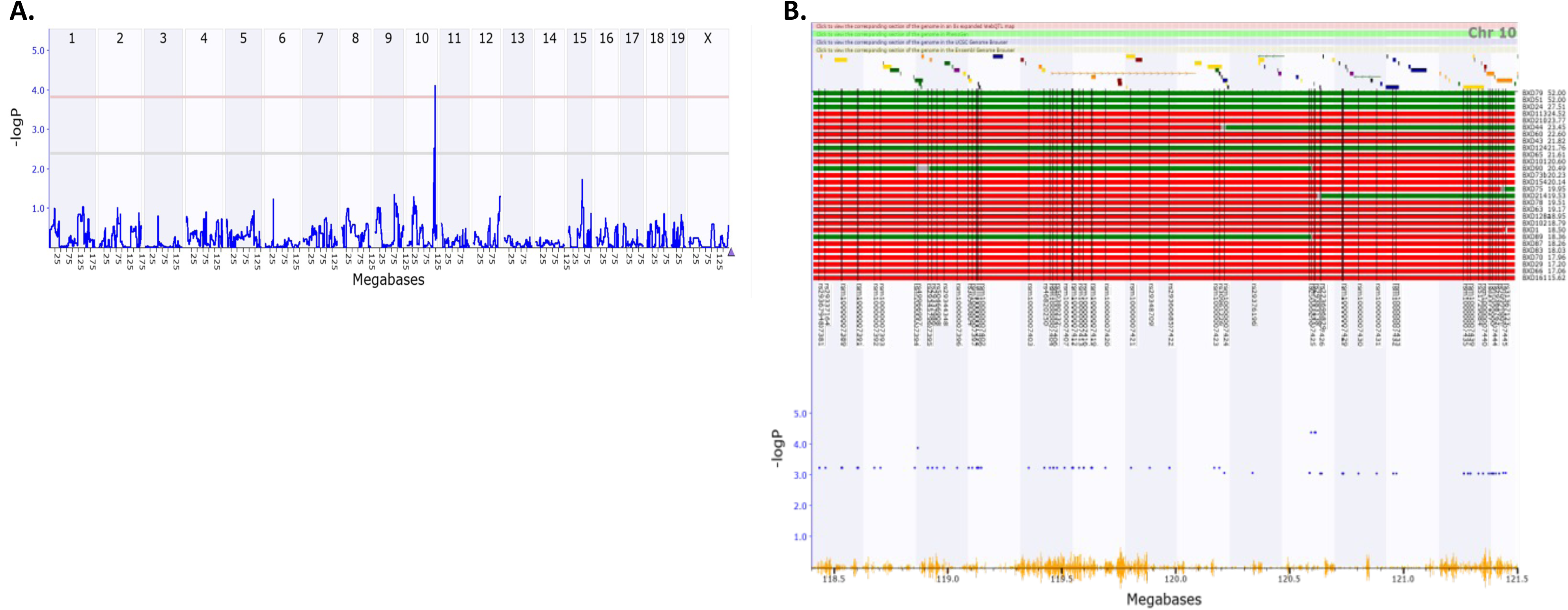
A significant quantitative trait locus (QTL) for tumor latency was identified on chromosome 10. QTLs associated with tumor latency (age in weeks) were examined in R/qtl2 and GEMMA. **A.** Genome wide QTL plot demonstrates the logarithm of the odds (LOD score (blue line) at each marker across the genome with chromosomes noted on the X – axis. The genome-wide significant threshold is indicated by pink line (-logP = 3.83). The genome-wide suggestive threshold of –logP = 2.39 is indicated by grey line. **B**. Zoomed image of the 1.5 LOD-drop confidence interval for the QTL on chromosome 10 is shown with megabases indicated on X axis. Red and green horizontal bars at the top of the figure represent the haplotypes at the position – green in the D-like DBA/2J-derived haplotype and red is the B-like C57BL/6J-derived haplotype. BXD-BC strains are aligned on top left with # indicating mean weeks. On the Y axis, blue dots represent the –log(p) linkage between a marker and tumor latency. Segregating SNPs in the BXD family are shown by the orange Seismograph at bottom.

**Table 4.** Chromosome 10 QTL Protein Coding Genes. An interval generated in GEMMA for chr10 (117.295-121.534 Mb) contains 121 annotated genes and open reading frames (ORFs), of which 27 were protein coding with gene symbol, description, start point, length, and variant or single nucleotide polymorphims (SNP) counts and density reported.

| Symbol | Gene Description | Mb Start | Length (Kb) | SNP Count | SNP Density |
| --- | --- | --- | --- | --- | --- |
| *Cpsf6* | cleavage and polyadenylation specific factor 6 | 117.344667 | 32.33 | 91 | 2.81 |
| *Cpm* | carboxypeptidase M | 117.6295 | 57.85 | 17 | 0.29 |
| *Mdm2* | transformed mouse 3T3 cell double minute 2 | 117.688875 | 21.88 | 7 | 0.32 |
| *Slc35e3* | solute carrier family 35, member E3 | 117.733678 | 12.68 | 1 | 0.08 |
| *Nup107* | nucleoporin 107 | 117.750621 | 42.12 | 35 | 0.83 |
| *Rap1b* | RAS related protein 1b | 117.814597 | 31.38 | 9 | 0.29 |
| *Mdm1* | transformed mouse 3T3 cell double minute 1 | 118.14154 | 27.46 | 68 | 2.48 |
| *Il22* | interleukin 22 | 118.204584 | 5.46 | 0 | 0.00 |
| *Il22b* | interleukin 22B | 118.289629 | 5.41 | 1 | 0.18 |
| *Ifng* | interferon gamma | 118.441046 | 4.85 | 18 | 3.71 |
| *Ifngas1* | Ifng antisense RNA 1 | 118.502035 | 54.49 | 59 | 1.08 |
| *Dyrk2* | dual-specificity tyrosine-(Y)-phosphorylation regulated kinase 2 | 118.855602 | 33.64 | 29 | 0.86 |
| *Cand1* | cullin associated and neddylation disassociated 1 | 119.198812 | 41.24 | 10 | 0.24 |
| *Grip1* | glutamate receptor interacting protein 1 | 119.453238 | 634.03 | 1711 | 2.70 |
| *Grip1os2* | glutamate receptor interacting protein 1, opposite strand 2 | 119.746062 | 15.64 | 130 | 8.31 |
| *Grip1os1* | glutamate receptor interacting protein 1, opposite strand 1 | 119.825204 | 27.19 | 114 | 4.19 |
| *Helb* | helicase (DNA) B | 120.083608 | 29.40 | 29 | 0.99 |
| *Irak3* | interleukin-1 receptor-associated kinase 3 | 120.141648 | 60.48 | 54 | 0.89 |
| *Tmbim4* | transmembrane BAX inhibitor motif containing 4 | 120.208826 | 16.07 | 3 | 0.19 |
| *Llph* | LLP homolog, long-term synaptic facilitation (Aplysia) | 120.227048 | 5.02 | 0 | 0.00 |
| *Hmga2* | high mobility group AT-hook 2 | 120.361268 | 115.20 | 56 | 0.49 |
| *Msrb3* | methionine sulfoxide reductase B3 | 120.7811 | 117.91 | 95 | 0.81 |
| *Lemd3* | LEM domain containing 3 | 120.923411 | 55.99 | 10 | 0.18 |
| *Wif1* | Wnt inhibitory factor 1 | 121.034004 | 66.64 | 18 | 0.27 |
| *Tbc1d30* | TBC1 domain family, member 30 | 121.263819 | 87.83 | 153 | 1.74 |
| *Gns* | glucosamine (N-acetyl)-6-sulfatase | 121.36509 | 32.16 | 111 | 3.45 |
| *Rassf3* | Ras association (RalGDS/AF-6) domain family member 3 | 121.41035 | 65.90 | 207 | 3.14 |

### Gene Variants and Expression QTL (eQTL) Analysis Identified Candidate Genes associated with Two QTLs and Histologic Traits

For necrosis score (BXD_27515) we identified a QTL on chr4 from 103.640707-105.243637 Mb with a –logP of 5.85 (**Supplemental Figs. 1A-B**). The QTL on chr4 associated with necrosis in the pathology analysis contained 23 predicted genes, of which only 6 are protein coding (*C8a*, *C8b*, *Dab1*, *Fyb2*, *Plpp3*, and *Prkaa2*) **Supplemental Table 6**. There are 3219 segregating variants in the region. No protein coding genes within the region had significant cis-eQTLs (*data not shown*).

For stroma and vascularity score (GN BXD_27514), we identified a QTL on chr12 from 64.387817-69.575599 Mb with a –logP of 5.12 (**Supplemental Figs. 1C-D**). The QTL on chr12 associated with vascularity and stroma score contained 172 predicted genes, of which 43 are annotated as protein coding **Supplemental Table** 7 There are 3750 annotated sequence variants which segregate. No protein coding genes within the region had significant cis-eQTLs (*data not shown*).

### Cross-species human phenome wide association study (Phe-WAS) comparisons reveal syntenic regions of interest

Once candidates were identified from the BXD-TNBC model, the translational validity of mouse candidate genes was tested. The human genomic regions syntenic to the significant mouse QTL regions identified on chromosome 10 for tumor latency and chromosomes 5 and 16 for tumor multiplicity were identified. For each QTL, the most proximal and distal genes were identified and the location of these genes in mouse and human were noted (**Table 5**). Human GWAS summary statistics available online from existing studies were used to carry out phenome wide association analysis (Phe-WAS) to examine regions of interest and the genes within them ^44–50^.

**Table 5.** Cross-species human phenome wide association study (Phe-WAS) identified genes and regions of interest.

|  |  |  |  |  |  |  |  |  |
| --- | --- | --- | --- | --- | --- | --- | --- | --- |
|  | Mouse proximal gene | Mouse distal gene | Mouse proximal gene location | Mouse distal gene location | Approx. mouse region length | Human proximal gene location | Human distal gene location | Approx. human region length |
| chr5 | *Spink2* | *Prdm8* | 77.16 | 98.24 | 21.08 | chr4:56.809 | chr4:80.204 | 23.395 |
| chr10 | *Cpsf6* | *Rassf3* | 117.358 | 121.41 | 4.051 | chr12:69.275 | chr12:64.61 | 4.665 |
| chr16 | *Heg1* | *Golgb1* | 33.684 | 36.885 | 3.201 | chr3:125.055 | chr3:121.663 | 3.392 |

## DISCUSSION

TNBC has limited personalized medicine approaches, restricting treatment primarily to conventional chemotherapy because TNBC is negative for classic targets including estrogen receptor (ER), progesterone receptor (PR), and amplification of the human epidermal growth factor receptor 2 (HER2). Due to these therapeutic limitations, patients with TNBC experience elevated recurrence and metastasis and poorer overall survival relative to patients with other BC subtypes ^54^. Further, TNBC disproportionately affects young and minority women including African American and Hispanic women which creates an urgency to correct this unmet clinical need and address these health disparities ^55^. Understanding specific underlying genetic modifiers and molecular mechanisms that drive TNBC tumor aggressiveness will improve our understanding and treatment of the disease.

Using our expertise in mouse model systems genetics, this study was designed to determine the interaction of modifier and causal genes in governing the heterogeneity of BC phenotypes. While previous work by our group using the C3Tag model to study pregnancy, obesity, overweight, and weight loss impacts on pre-neoplastic lesions (hyperplasia and DCIS) as well as cancer burden, progression, and response to therapy ^9–13^, limitations around GEMMs in general drove us to create a more robust model. Current BC GEMMs lack genetic complexity because mice are on a single inbred background which impairs the rigorous investigation into individual genetic variation and tumor initiation, progression, or response to therapy^56–58^. Because of this limitation, pre-clinical models typically fail to translate well to impact patient care.

Herein, we created a model system, the BXD-BC, which demonstrate large, heritable variation in multiple breast tumor phenotypes. Importantly, the parent FVB strain for C3Tag GEMMs has above average latency and burden, but lower survival and multiplicity compared to novel BXD-TNBC strains; these results demonstrate that crosses into the BXD strains modulate susceptibility. Because the BXD intercross developed BXD-BC mouse strains with demonstrated *accelerated* tumor onset (i.e. mice with younger latency) or *delayed* tumor onset (i.e. older latency), using extremes of latency in future studies may aid in identifying the molecular underpinnings of pathways such as DNA repair or genomic instability in this model. Previous successes using the BXD family with a systems genetics approach allowed the identification of a novel, clinically relevant, and drug-responsive gene product for intraocular pressure now in trials ^59,60^.

Heritability (h2) and significance of strain calculations demonstrated extent of phenotypic variance due to germline genetics (i.e. strain) versus spontaneous somatic or environmental variation. High heritability increases the likelihood of identifying a significant QTL. The most significantly heritable traits were latency, endpoint, and multiplicity. When two strains that did not develop tumors out till 1 year (BXD-BC51 and BXD-BC79), heritability of survival was the greatest trait regulated. Why the strains failed to present with mammary tumors is of great interest. The strains were genotyped correctly and the expression in tumors of T antigen was confirmed. Therefore, ongoing studies including adding more progeny and strains around the variants using additional BXD strains associated with latency are planned.

There are several limitations to our current findings. Although estimated heritability values for many traits were high, only a few significant QTLs were identified due to a relatively small number of strains. Moreover, tumor traits are influenced by many genetic loci, as in tumor multiplicity. As more loci contribute to a trait, the effect size (as a proportion of total variance) of any single QTL decreases, reducing power to detect them. Therefore, increasing the number of strains to increase the power to detect QTL, and provide greater precision for the QTL already identified, is paramount for future studies.

Significant QTLs were identified based on traits including tumor multiplicity, latency, and histologic qualities and plausible candidate genes. For tumor multiplicity, several loci across chromosomes contributed to the trait when strains that never developed tumors were excluded. Four QTLs were first identified on Chr5, Chr8, Chr16, and Chr18, with the highest LOD being on chr16. When chr5 QTL was controlled for in remapping, the chr16 QTL became significant and strong, and vice versa. The three loci with the highest LOD scores (on chr5, chr12, chr16) were of the C57BL/6 (B) genotype at all three loci. Interestingly, the two strains that never develop tumors have the DBA/2J genotype at the chr5 and chr12 QTLs, and the B genotype at the chr16 QTL. These data suggest that several loci contribute to multiplicity in the BXD-BC model, yet the additive or interactive nature of loci remains unclear and is under investigation by increasing the number of strains tested. The chr16 QTL interval associated with tumor multiplicity identified candidate genes in with only 14 genes including variants predicted to alter protein function including *Ccdc14*, *Cd86*, *Dirc2*, *Dtx3l*, *Gm49662*, *Hspbap1*, *Mylk*, *Parp14*, *Parp9*, *Ptplb*, *Sema5b*, *Slc15a2*, *Slc49a4*, *Wdr5b*. Candidate genes were narrowed down by cis-eQTL analysis which identified 8 genes in this chr16 QTL loci including *Dirc2*, *Dtx3l*, *Stfa3*, and *Zdhhc23* in T helper or regulator T cells. Genes identified using spleen or white adipose tissue included *Kiaa0226*, *Hspbap1*, *Popdc2*, and *Slc15a2*. For tumor latency, a significant QTL was identified on Chr10 when the strains that never developed tumors were included in analysis due to the strong influence on heritability. Gene candidates identified using two approaches identified 6 protein coding candidates of interest *Msrb3*, *Lemd3*, *Wif1*, *Tbc1d30*, *Gns*, and *Rassf3*. Candidate genes were narrowed down by cis-eQTL analysis including *Cand1*, *Tbc1d30,* and *Rassf3* in tissues relevant to cancer. TBC1 domain family member 30 (*Tbc1d30* ^61,62^ and Glucosamine (N-Acetyl)-6-Sulfatase *(Gns* ^63^) are of particular interest due to missense and splice variants, respectively, that are precited to impact protein function.

*Rassf3* is a high likelihood candidate to modulate BXD-BC tumor traits – it is in a narrow QTL interval identified on chr10, has a significant *cis*-eQTL in a number of relevant tissues. *Rassf3* is part of the Ras association domain family (RASSF) gene family, and inactivation of these genes have been found in many human tumors^64,65^. *Rassf3*’s role in cancer is predicted to be through stabilizing p53^66^. The presence of *Rassf3* delays mammary tumor formation in a murine model^67^. Finding candidates such as *Rassf3* validates the BXD-BC approach.

Taken together, multiple QTLs and eQTLs were identified which yielded candidate genes of interest that associated with significantly heritable tumor traits. Future work is necessary to narrow down QTLs of interest and *in vitro* and *in vivo* work must be conducted to validate candidate genes in TNBC risk and progression, such as *Rassf3*. The challenge of precision medicine is to model complex interactions among DNA variants, sets of phenotypes, and diverse environmental factors and confounders. This challenge can be met by increasing the translatability and power of pre-clinical models. Ideally, outcomes from the BXD-BC GEMM tool presented herein and in future studies will be transformative with the identification of conserved, biologically relevant targets including genes and networks underlying individual differences in BC phenotypes and treatment responses, which advance our long-term goals to improve personalized medicine. This novel model combined with the rich BXD GeneNetwork database provides a unique and enduring BC model for future systems genetics analyses including gene-by-environment interactions. The results generated will fundamentally advance the field of BC genetics and enable targeted mechanistic studies to better understand risk and improve therapeutic outcomes for BC.

## Acknowledgments

We acknowledge support from the following funding sources:

National Cancer Institute (NCI) CA262112 (DNH, LL, RW, DGA, LM)

National Institutes of Health grant NCI R01CA253329 (LM)

UTHSC Center for Cancer Research (LM)

American Association for Cancer Research (AACR)-Triple Negative Breast Cancer Foundation Research Fellowship (LMS)

National Institutes of Health grant NCI F32 CA250192 (LMS)

National Institute of Health grant NCI F30CA265224 (JRH)

National Institute of Health grant NCI U24CA210988 (DNH)

National Institute of Health grant NCI UG1CA233333 (DNH)

## Acknowledgments

We thank Daniel Johnson from UTHSC Molecular Resource Center.

## Data Availability Statement

The data generated in this study are available within the article and its supplementary data files.

## Legends

**Supplemental Figure 1. A significant quantitative trait locus (QTL) for tumor histologic scores were identified on chromosomes 4 and 12**. QTLs associated with necrosis as scored by histology were examined in GEMMA. **A.** Genome wide QTL plot demonstrates the logarithm of the odds (LOD score (blue line) at each marker across the genome with chromosomes noted on the X – axis for tumor necrosis. The genome-wide significant threshold is indicated by pink line (– logP = 3.82). The genome-wide suggestive threshold of –logP = 2.94 is indicated by grey line. **B**. Zoomed image of the 1.5 LOD-drop confidence interval for the QTL on chromosome 4 is shown with megabases indicated on X axis. Red and green horizontal bars at the top of the figure represent the haplotypes at the position – green in the D-like DBA/2J-derived haplotype and red is the B-like C57BL/6J-derived haplotype. BXD-BC strains are aligned on top left with # indicating necrosis score. On the Y axis, blue dots represent the –log(p) linkage between a marker and tumor latency. Segregating SNPs in the BXD family are shown by the orange Seismograph at bottom. **C.** Genome wide QTL plot for histological score for stroma and vascularity, with genome-wide significant threshold is indicated by pink line (-logP = 3.69). The genome-wide suggestive threshold of –logP = 2.78 is indicated by grey line. **D**. Zoomed image of the 1.5 LOD-drop confidence interval for the QTL on chromosome 12 is shown with megabases indicated on X axis. BXD-BC strains are aligned on top left with # indicating histologic score for stroma and vascularity.

