## Supplementary figures and images for "Determining susceptibility loci in triple negative breast cancer using a novel pre-clinical model"

### Supplementary Figure 1

#### Supplemental Figure 1

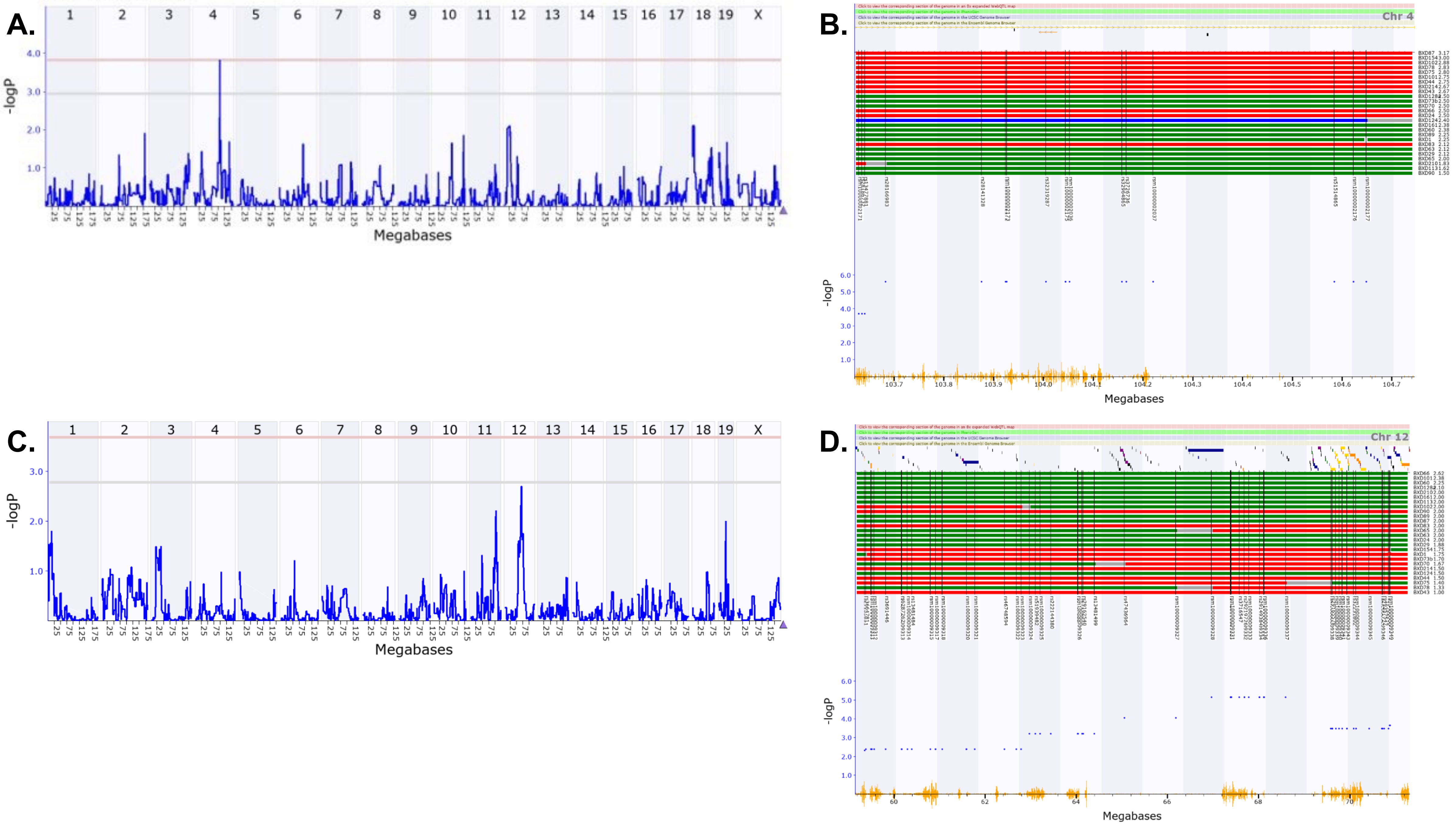
