## Supplementary Table 1 for "Determining susceptibility loci in triple negative breast cancer using a novel pre-clinical model"

**Supplemental Table 1. BXD-BC trait IDs based on tumor characteristics and histologic analysis entered into GeneNetwork.**

| BXD Record ID# | Phenotype, Description, and Units |
|---|---|
| 27516 | Cancer: Breast cancer inflammation, tumor elicited inflammation, 0 = none, 1= minimal, 2= mild, 3 moderate, 4 heavy, 5=extensive inflammation, in female F1 (female BXD crossed to male C3 (1)-TAg (JAX Stock 013591: FVB-Tg(C3-1TAg)cJeg/JegJ, SV40 T/t-antig |
| 21537 | Cancer: Breast cancer age at morbidity (euthanasia) in female F1 (female BXD crossed to male C3 (1)-TAg (JAX Stock 013591: FVB-Tg(C3-1TAg)cJeg/JegJ, SV40 T/t-antigen) strain mean including never developed [age in weeks] |
| 24401 | Cancer: Breast cancer age at morbidity (euthanasia) in female F1 (female BXD crossed to male C3 (1)-TAg (JAX Stock 013591: FVB-Tg(C3-1TAg)cJeg/JegJ, SV40 T/t-antigen) strain mean including strains that never developed tumors [age in days] |
| 21527 | Cancer: Breast cancer age at morbidity (euthanasia) in female F1 (female BXD crossed to male C3 (1)-TAg (JAX Stock 013591: FVB-Tg(C3-1TAg)cJeg/JegJ, SV40 T/t-antigen) strain mean [age in days] |
| 27513 | Cancer: Breast cancer Epithelial-mesenchymal transition (EMT) on 5-point scale, 0 = no pleomorphism, 1 = subepithelial pleomorphism, 2 = faint streaming, 3 = distinct frequent streaming, 4 = distinct sarcomatous transition seen, in female F1 (female BXD c |
| 21529 | Cancer: Breast cancer Mean volume of first tumour in female F1 (female BXD crossed to male C3 (1)-TAg (JAX Stock 013591: FVB-Tg(C3-1TAg)cJeg/JegJ, SV40 T/t-antigen) strain mean (mm^3). |
| 27517 | Cancer: Breast cancer mitosis, as number of mitoses per high-power field (40X) averaged across 3 or more areas of tumor in female F1 (female BXD crossed to male C3 (1)-TAg (JAX Stock 013591: FVB-Tg(C3-1TAg)cJeg/JegJ, SV40 T/t-antigen) strain mean |
| 27515 | Cancer: Breast cancer necrosis, prevalence of necrosis and cavitary tumor loss, 0 = none, 1= minimal, 2= mild, 3 moderate, 4 heavy, 5=extensive necrosis, in female F1 (female BXD crossed to male C3 (1)-TAg (JAX Stock 013591: FVB-Tg(C3-1TAg)cJeg/JegJ, SV40 |
| 21528 | Cancer: Breast cancer number of tumors at morbidity (euthanasia) in female F1 (female BXD crossed to male C3 (1)-TAg (JAX Stock 013591: FVB-Tg(C3-1TAg)cJeg/JegJ, SV40 T/t-antigen) Strain mean (n) |
| 24403 | Cancer: Breast cancer number of tumors at morbidity (euthanasia) in female F1 (female BXD crossed to male C3 (1)-TAg (JAX Stock 013591: FVB-Tg(C3-1TAg)cJeg/JegJ, SV40 T/t-antigen) strain mean including strains that never developed tumors (n) |
| 21530 | Cancer: Breast cancer total tumor volume at morbidity (euthanasia) in female F1 (female BXD crossed to male C3 (1)-TAg (JAX Stock 013591: FVB-Tg(C3-1TAg)cJeg/JegJ, SV40 T/t-antigen) -Strain mean (mm^3) |
| 24405 | Cancer: Breast cancer total tumor volume at morbidity (euthanasia) in female F1 (female BXD crossed to male C3 (1)-TAg (JAX Stock 013591: FVB-Tg(C3-1TAg)cJeg/JegJ, SV40 T/t-antigen) strain mean including strains that never developed tumors with strains th |
| 21532 | Cancer: Breast cancer total weight of tumours at morbidity (euthanasia) in female F1 (female BXD crossed to male C3 (1)-TAg (JAX Stock 013591: FVB-Tg(C3-1TAg)cJeg/JegJ, SV40 T/t-antigen) - strain mean (mg) |
| 24407 | Cancer: Breast cancer total weight of tumours at morbidity (euthanasia) in female F1 (female BXD crossed to male C3 (1)-TAg (JAX Stock 013591: FVB-Tg(C3-1TAg)cJeg/JegJ, SV40 T/t-antigen) strain mean including strains that never developed tumors as 0 (mg) |
| 24402 | Cancer: Breast cancer tumor development latency in female F1 (female BXD crossed to male C3 (1)-TAg (JAX Stock 013591: FVB-Tg(C3-1TAg)cJeg/JegJ, SV40 T/t-antigen) strain mean including strains that never developed tumors as 52 weeks [age in weeks] |
| 21526 | Cancer: Breast cancer tumor development latency in female F1 (female BXD crossed to male C3 (1)-TAg (JAX Stock 013591: FVB-Tg(C3-1TAg)cJeg/JegJ, SV40 T/t-antigen) strain mean [age in weeks] |
| 24412 | Cancer: Breast cancer tumor time between first detection and euthanasia in female F1 (female BXD crossed to male C3 (1)-TAg (JAX Stock 013591: FVB-Tg(C3-1TAg)cJeg/JegJ, SV40 T/t-antigen) strain mean including strains that never developed tumors [days] |
| 24408 | Cancer: Breast cancer tumor, one tumor greater than 2cm at death in female F1 (female BXD crossed to male C3 (1)-TAg (JAX Stock 013591: FVB-Tg(C3-1TAg)cJeg/JegJ, SV40 T/t-antigen) strain mean including strains that never developed tumors as 0 [n] |
| 24398 | Cancer: Breast cancer tumor, one tumor greater than 2cm at morbidity (euthanasia) in female F1 (female BXD crossed to male C3 (1)-TAg (JAX Stock 013591: FVB-Tg(C3-1TAg)cJeg/JegJ, SV40 T/t-antigen). Strain mean [n] |
| 27514 | Cancer: Breast cancer vascularity and stroma score, 0 = none, 1 = minimal stroma, 2= mild, 3 = moderate, 4 = heavy, 5 = scirrhous, in female F1 (female BXD crossed to male C3 (1)-TAg (JAX Stock 013591: FVB-Tg(C3-1TAg)cJeg/JegJ, SV40 T/t-antigen) strain me |
| 24404 | Cancer: Breast cancer volume of first tumor in female F1 (female BXD crossed to male C3 (1)-TAg (JAX Stock 013591: FVB-Tg(C3-1TAg)cJeg/JegJ, SV40 T/t-antigen) strain strain mean including strains that never developed tumors with strains that never develop |
| 21531 | Cancer: Breast cancer weight of first tumour at morbidity (euthanasia) in female F1 (female BXD crossed to male C3 (1)-TAg (JAX Stock 013591: FVB-Tg(C3-1TAg)cJeg/JegJ, SV40 T/t-antigen) - strain mean (mg) |
| 24406 | Cancer: Breast cancer weight of first tumour at morbidity (euthanasia) in female F1 (female BXD crossed to male C3 (1)-TAg (JAX Stock 013591: FVB-Tg(C3-1TAg)cJeg/JegJ, SV40 T/t-antigen) strain mean including strains that never developed tumors as 0 (mg) |
