## Supplementary Table 2 for "Determining susceptibility loci in triple negative breast cancer using a novel pre-clinical model"

**Supplemental Table 2. Multiple Tumor Traits Display Significant Heritability in BXD-BC Strains.** Heritability (h2) and significance of strain effect (p) are shown tumor traits collected for N=28 BXD-BC strains, with an average of 8 replicates per strain. Strain effect was tested by ANOVA. Strains that did not develop tumors (BXD-BC51 and BXD-BC79) were included. Bold indicates significance P<0.05.

| Phenotype (Trait) | h2 | p-value |
|---|---|---|
| Survival (Time from latency to endpoint) | 0.970 | **1.48E-134** |
| Age at endpoint | 0.926 | **8.90E-97** |
| Tumor latency | 0.524 | **9.63E-19** |
| Multiplicity | 0.483 | **1.38E-16** |
| Total tumor weight | 0.317 | **6.42E-07** |
| Total tumor volume | 0.303 | **2.74E-06** |
| Endpoint 1 tumor >2cm | 0.203 | **0.011** |
| Endpoint 3 tumors >1cm | 0.197 | **0.017** |
| T1 tumor weight | 0.175 | 0.060 |
| T1 volume | 0.170 | 0.078 |
| T1 tumor location | 0.138 | 0.359 |
