## Supplementary Table 3 for "Determining susceptibility loci in triple negative breast cancer using a novel pre-clinical model"

**Supplemental Table 3. Tumor Histology Traits Display Insignificant Heritability in all BXD-BC Strains.** Heritability (h2) and significance of strain effect (p) are shown tumor traits collected for N=26 BXD-BC strains, with an average of 8 replicates per strain. Strain effect was tested by ANOVA. Strains that did not develop tumors were not included. Scored histology was averaged from over 3 randomly selected regions of interest in tumor H+E section.

| Phenotype (Trait) | h2 | p-value |
|---|---|---|
| Mitosis per high powered field (40X) | 0.34 | 0.097 |
| Vascularity and stroma | 0.33 | 0.101 |
| Necrosis | 0.26 | 0.44 |
| Epithelial Mesenchymal Transition (EMT) | 0.24 | 0.557 |
