## Supplementary Table 4 for "Determining susceptibility loci in triple negative breast cancer using a novel pre-clinical model"

**Supplemental Table 4. Chromosome 16 eQTL associated with latency.** Examination genes identified within Chr16 by cis-eQTLs across tissues relevant to tumor or immune biology, including leucocytes, neutrophils, spleen, white adipose tissue, T-helper cells, and regulatory T cells. r and rho correlations are indicated with associated P values. P values <0.05 are highlighted bold and grey.

| Gene | Tissue (mRNA) | Links | Probe ID | .-logP | Peak Position | TUMOR MULTIPLICITY_Strains including without tumor (BXD_ 24403) | | | | TUMOR MULTIPLICITY_Strains with only tumor (BXD_ 21528) | | | |
|---|---|---|---|---|---|---|---|---|---|---|---|---|---|
| | | | | | | **r** | **r p-value** | **rho** | **rho p-value** | **r** | **r p-value** | **rho** | **rho p-value** |
| *Hspbap1* | Leucocytes | https://genenetwork.org/show_trait?trait_id=ILM4060022&dataset=Illum_BXD_PBL_1108 | ILM4060022 | 2.5 | N/a | -0.101 | 0.899 | -0.4 | 0.6 | -0.993 | 0.076 | -1 | **9.00E-11** |
| *Hspbap1* | Neutrophils | https://genenetwork.org/show_trait?trait_id=17325261&dataset=UTHSC-Neut-1014 | 17325261 | 4.7 | Chr16: 35.770386 | 0.005 | 0.991 | 0.119 | 0.779 | 0.073 | 0.876 | 0.036 | 0.939 |
| *Hspbap1* | Spleen | https://genenetwork.org/show_trait?trait_id=10435443&dataset=UTHSC_SPL_RMA_1210 | 10435443 | 6.9 | Chr16: 35.770407 | -0.047 | 0.841 | -0.162 | 0.484 | -0.162 | 0.508 | -0.24 | 0.322 |
| *Hspbap1* | White Adipose Tissue | https://genenetwork.org/show_trait?trait_id=17325261&dataset=EPFLADGL1013 | 17325261 | 1.7 | Chr16: 35.770386 | -0.162 | 0.549 | -0.175 | 0.517 | -0.478 | 0.084 | -0.367 | 0.197 |
| *Hspbap1* | T-helper Cells | https://genenetwork.org/show_trait?trait_id=1428566_at&dataset=RTHC_0211_R | 1428566_at | 4.0 | Chr16: 35.824782 | -0.093 | 0.861 | -0.314 | **-0.314** | -0.807 | 0.099 | -0.8 | 0.104 |
| *Hspbap1* | T-regulatory Cells | https://genenetwork.org/show_trait?trait_id=1428566_at&dataset=RTC_1106_R | 1428566_at | 8.2 | Chr16: 35.824782 | 0.407 | 0.423 | -0.029 | 0.957 | -0.391 | 0.515 | -0.8 | 0.104 |
| *Slc15a2* | Leucocytes | https://genenetwork.org/show_trait?trait_id=ILM1780538&dataset=Illum_BXD_PBL_1108 | ILM1780538 | 9.8 | Chr16: 36.750469 | 0.346 | 0.654 | 0.6 | 0.4 | 0.285 | 0.816 | 0.5 | 0.667 |
| *Slc15a2* | Neutrophils | https://genenetwork.org/show_trait?trait_id=17330218&dataset=UTHSC-Neut-1014 | 17330218 | 8.4 | Chr16: 36.750164 | -0.392 | 0.336 | -0.024 | 0.955 | 0.037 | 0.937 | 0.357 | 0.432 |
| *Slc15a2* | Spleen | https://genenetwork.org/show_trait?trait_id=10439321&dataset=UTHSC_SPL_RMA_1210 | 10439321 | 8.2 | Chr16: 36.750181 | -0.264 | 0.247 | -0.153 | 0.507 | -0.158 | 0.518 | -0.084 | 0.732 |
| *Slc15a2* | White Adipose Tissue | https://genenetwork.org/show_trait?trait_id=17330218&dataset=EPFLADGL1013 | 17330218 | 9.0 | Chr16: 36.750164 | -0.203 | 0.45 | -0.04 | 0.884 | 0.098 | 0.739 | 0.213 | 0.464 |
| *Slc15a2* | T-helper Cells | https://genenetwork.org/show_trait?trait_id=1417600_at&dataset=RTHC_0211_R | 1417600_at | 10.2 | Chr16: 36.750199 | -0.301 | 0.563 | -0.086 | 0.872 | 0.484 | 0.409 | 0.6 | 0.285 |
| *Slc15a2* | T-regulatory Cells | https://genenetwork.org/show_trait?trait_id=1417600_at&dataset=RTC_1106_R | 1417600_at | 9.7 | Chr16: 36.750199 | -0.236 | 0.652 | -0.086 | 0.872 | 0.516 | 0.373 | 0.6 | 0.285 |
| *Kiaa0226* | Leucocytes | N/a | N/a | N/a | N/a | N/a | N/a | N/a | N/a | N/a | N/a | N/a | N/a |
| *Kiaa0226* | Neutrophils | https://genenetwork.org/show_trait?trait_id=17329877&dataset=UTHSC-Neut-1014 | 17329877 | 9.6 | Chr16: 32.821702 | -0.141 | 0.739 | -0.238 | 0.57 | -0.412 | 0.358 | -0.286 | 0.535 |
| *Kiaa0226* | Spleen | https://genenetwork.org/show_trait?trait_id=10439092&dataset=UTHSC_SPL_RMA_1210 | 10439092 | 10.1 | Chr16: 32.821707 | 0.23 | 0.315 | 0.362 | 0.107 | 0.515 | **0.024** | 0.54 | **0.017** |
| *Kiaa0226* | White Adipose Tissue | N/a | N/a | N/a | N/a | N/a | N/a | N/a | N/a | N/a | N/a | N/a | N/a |
| *Kiaa0226* | T-helper Cells | N/a | N/a | N/a | N/a | N/a | N/a | N/a | N/a | N/a | N/a | N/a | N/a |
| *Kiaa0226* | T-regulatory Cells | N/a | N/a | N/a | N/a | N/a | N/a | N/a | N/a | N/a | N/a | N/a | N/a |
| *Popdc2* | Leucocytes | https://genenetwork.org/show_trait?trait_id=ILM2030056&dataset=Illum_BXD_PBL_1108 | ILM2030056 | 6.9 | Chr16: 38.377934 | 0.201 | 0.799 | 0.4 | 0.6 | -0.924 | 0.251 | -0.5 | 0.667 |
| *Popdc2* | Neutrophils | https://genenetwork.org/show_trait?trait_id=17325592&dataset=UTHSC-Neut-1014 | 17325592 | 7.2 | Chr16: 38.362173 | -0.65 | 0.081 | -0.611 | 0.108 | -0.503 | 0.25 | -0.523 | 0.229 |
| *Popdc2* | Spleen | https://genenetwork.org/show_trait?trait_id=10435697&dataset=UTHSC_SPL_RMA_1210 | 10435697 | 8.0 | Chr16: 38.362209 | 0.065 | 0.779 | -0.12 | 0.606 | -0.116 | 0.635 | -0.275 | 0.254 |
| *Popdc2* | White Adipose Tissue | https://genenetwork.org/show_trait?trait_id=17325592&dataset=EPFLADGL1013 | 17325592 | 9.5 | Chr16: 38.362173 | 0.514 | **0.042** | 0.284 | 0.286 | 0.341 | 0.233 | 0.095 | 0.748 |
| *Popdc2* | T-helper Cells | https://genenetwork.org/show_trait?trait_id=1417806_at&dataset=RTHC_0211_R | 1417806_at | 3.3 | Chr16: 38.377659 | 0.235 | 0.654 | -0.143 | 0.787 | -0.207 | 0.739 | -0.7 | 0.188 |
| *Popdc2* | T-regulatory Cells | N/a | N/a | N/a | N/a | N/a | N/a | N/a | N/a | N/a | N/a | N/a | N/a |
| *Dirc2* | Leucocytes | https://genenetwork.org/show_trait?trait_id=ILM2810494&dataset=Illum_BXD_PBL_1108 | ILM2810494 | 3.0 | Chr16: 35.695101 | 0.322 | 0.678 | 0 | 1 | -0.163 | 0.895 | -0.5 | 0.667 |
| *Dirc2* | Neutrophils | https://genenetwork.org/show_trait?trait_id=17330080&dataset=UTHSC-Neut-1014 | 17330080 | 2.1 | Chr16: 35.694062 | 0.071 | 0.867 | 0.381 | 0.352 | 0.452 | 0.309 | 0.607 | 0.148 |
| *Dirc2* | Spleen | https://genenetwork.org/show_trait?trait_id=10439239&dataset=UTHSC_SPL_RMA_1210 | 10439239 | 3.7 | Chr16: 35.694903 | 0.47 | **0.031** | 0.376 | 0.093 | 0.423 | 0.071 | 0.246 | 0.311 |
| *Dirc2* | White Adipose Tissue | https://genenetwork.org/show_trait?trait_id=17330080&dataset=EPFLADGL1013 | 17330080 | 2.2 | Chr16: 35.694062 | 0.111 | 0.684 | -0.165 | 0.542 | 0.016 | 0.957 | -0.284 | 0.326 |
| *Dirc2* | T-helper Cells | https://genenetwork.org/show_trait?trait_id=1454654_at&dataset=RTHC_0211_R | 1454654_at | 3.0 | Chr16: 35.697399 | -0.461 | 0.358 | -0.029 | 0.957 | 0.283 | 0.644 | 0.7 | 0.188 |
| *Dirc2* | T-regulatory Cells | https://genenetwork.org/show_trait?trait_id=1454654_at&dataset=RTC_1106_R | 1454654_at | 2.3 | Chr16: 35.697399 | -0.679 | 0.138 | -0.543 | 0.266 | 0.179 | 0.773 | -0.2 | 0.747 |
| *Dtx3l* | Leucocytes | https://genenetwork.org/show_trait?trait_id=ILM1050168&dataset=Illum_BXD_PBL_1108 | ILM1050168 | 2.6 | Chr16: 35.928239 | -0.629 | 0.371 | -1 | **1.00E-20** | -0.932 | 0.237 | -1 | **9.00E-11** |
| *Dtx3l* | Neutrophils | https://genenetwork.org/show_trait?trait_id=17330119&dataset=UTHSC-Neut-1014 | 17330119 | 2.3 | Chr16: 35.926511 | -0.051 | 0.905 | -0.119 | 0.779 | -0.554 | 0.197 | -0.679 | 0.094 |
| *Dtx3l* | Spleen | https://genenetwork.org/show_trait?trait_id=10439268&dataset=UTHSC_SPL_RMA_1210 | 10439268 | 3.4 | Chr16: 35.926511 | -0.256 | 0.263 | -0.066 | 0.777 | 0.06 | 0.806 | 0.188 | 0.441 |
| *Dtx3l* | White Adipose Tissue | https://genenetwork.org/show_trait?trait_id=17330119&dataset=EPFLADGL1013 | 17330119 | 3.4 | Chr16: 35.926511 | -0.34 | 0.197 | -0.49 | 0.054 | -0.175 | 0.55 | -0.424 | 0.131 |
| *Dtx3l* | T-helper Cells | https://genenetwork.org/show_trait?trait_id=1435208_at&dataset=RTHC_0211_R | 1435208_at | 3.1 | Chr16: 35.926596 | -0.345 | 0.503 | 0.086 | 0.872 | 0.644 | 0.241 | 0.9 | **0.037** |
| *Dtx3l* | T-regulatory Cells | https://genenetwork.org/show_trait?trait_id=1435208_at&dataset=RTC_1106_R | 1435208_at | 3.4 | Chr16: 35.926596 | -0.224 | 0.67 | 0.086 | 0.872 | 0.705 | 0.183 | 0.9 | **0.037** |
| *Stfa3* | Leucocytes | https://genenetwork.org/show_trait?trait_id=ILM130504&dataset=Illum_BXD_PBL_1108 | ILM130504 | 1.5 | Chr16: 36.452150 | -0.574 | 0.426 | -0.8 | 0.2 | -0.457 | 0.698 | -0.5 | 0.667 |
| *Stfa3* | Neutrophils | https://genenetwork.org/show_trait?trait_id=17330183&dataset=UTHSC-Neut-1014 | 17330183 | 4.2 | Chr16: 36.450537 | 0.115 | 0.786 | 0.024 | 0.955 | -0.13 | 0.781 | -0.143 | 0.76 |
| *Stfa3* | Spleen | https://genenetwork.org/show_trait?trait_id=10439299&dataset=UTHSC_SPL_RMA_1210 | 10439299 | 1.9 | Chr16: 35.171449 | 0.024 | 0.916 | -0.021 | 0.929 | -0.024 | 0.923 | -0.039 | 0.875 |
| *Stfa3* | White Adipose Tissue | https://genenetwork.org/show_trait?trait_id=17330183&dataset=EPFLADGL1013 | 17330183 | 2.6 | Chr16: 36.450537 | -0.293 | 0.27 | -0.299 | 0.261 | 0.087 | 0.768 | -0.075 | 0.799 |
| *Stfa3* | T-helper Cells | https://genenetwork.org/show_trait?trait_id=1419709_at&dataset=RTHC_0211_R | 1419709_at | 5.9 | Chr16: 36.450554 | 0.299 | 0.566 | -0.029 | 0.957 | -0.567 | 0.319 | -0.8 | 0.104 |
| *Stfa3* | T-regulatory Cells | https://genenetwork.org/show_trait?trait_id=1419709_at&dataset=RTC_1106_R | 1419709_at | 5.7 | Chr16: 36.450554 | 0.407 | 0.424 | 0.086 | 0.872 | -0.23 | 0.709 | -0.6 | 0.285 |
| *Zdhhc23* | Leucocytes | https://genenetwork.org/show_trait?trait_id=ILM4390196&dataset=Illum_BXD_PBL_1108 | ILM4390196 | 3.0 | Chr16: 43.888484 | -0.869 | 0.131 | -0.4 | 0.6 | 0.83 | 0.377 | 0.5 | 0.667 |
| *Zdhhc23* | Neutrophils | https://genenetwork.org/show_trait?trait_id=17330499&dataset=UTHSC-Neut-1014 | 17330499 | 2.2 | Chr16: 43.969078 | 0.264 | 0.527 | 0.024 | 0.955 | -0.671 | 0.099 | -0.464 | 0.294 |
| *Zdhhc23* | Spleen | https://genenetwork.org/show_trait?trait_id=10439542&dataset=UTHSC_SPL_RMA_1210 | 10439542 | 4.4 | Chr16: 43.969146 | 0.294 | 0.195 | 0.15 | 0.516 | 0.059 | 0.81 | -0.011 | 0.966 |
| *Zdhhc23* | White Adipose Tissue | https://genenetwork.org/show_trait?trait_id=17330499&dataset=EPFLADGL1013 | 17330499 | 4.3 | Chr16: 43.969078 | 0.233 | 0.385 | 0.374 | 0.154 | 0.171 | 0.558 | 0.341 | 0.233 |
| *Zdhhc23* | T-helper Cells | https://genenetwork.org/show_trait?trait_id=1441069_at&dataset=RTHC_0211_R | 1441069_at | 5.9 | Chr16: 43.965083 | -0.23 | 0.661 | -0.371 | 0.468 | -0.873 | 0.053 | -0.9 | **0.037** |
| *Zdhhc23* | T-regulatory Cells | https://genenetwork.org/show_trait?trait_id=1441069_at&dataset=RTC_1106_R | 1441069_at | 5.4 | Chr16: 43.965083 | N/a | N/a | N/a | N/a | N/a | N/a | N/a | N/a |
