## Supplementary Table 5 for "Determining susceptibility loci in triple negative breast cancer using a novel pre-clinical model"

**Supplemental Table 5. Chromosome 10 eQTL associated with latency.** Examination genes identified within Chr10 by cis-eQTLs across tissues relevant to tumor or immune biology, including leucocytes, neutrophils, spleen, white adipose tissue, T-helper cells, and regulatory T cells. r and rho correlations are indicated with associated P values. P values <0.05 are highlighted bold and grey.

| Gene | Tissue (mRNA) | Links | Probe ID | .-logP | Peak Position | TUMOR LATENCY_strains including without tumor (BXD_ 24402) | | | | TUMOR MULTIPLICITY_Strains including without tumor (BXD_ 24403) | | | | TUMOR LATENCY_strains with only tumor (BXD_ 21526) | | | | TUMOR MULTIPLICITY_Strains with only tumor (BXD_ 21528) | | | |
|---|---|---|---|---|---|---|---|---|---|---|---|---|---|---|---|---|---|---|---|---|---|
| | | | | | | r | r p-value | rho | rho p-value | r | r p-value | rho | rho p-value | r | r p-value | rho | rho p-value | r | r p-value | rho | rho p-value |
| *Rassf3* | Leucocytes | https://genenetwork.org/show_trait?trait_id=ILM730300&dataset=Illum_BXD_PBL_1108 | ILM730300 | 2.4 | Chr10: 121.410559 | -0.875 | 0.125 | -1 | **1.00E-20** | 0.79 | 0.210 | 0.8 | 0.200 | -0.998 | **0.045** | -1 | **9.00E-11** | 0.749 | 0.461 | 0.5 | 0.667 |
| *Rassf3* | Neutrophils | https://genenetwork.org/show_trait?trait_id=17245481&dataset=UTHSC-Neut-1014 | 17245481 | 3.2 | Chr10: 121.410350 | -0.026 | 0.951 | -0.333 | 0.420 | -0.009 | 0.983 | -0.048 | 0.911 | 0.001 | 0.998 | -0.286 | 0.535 | -0.043 | 0.927 | -0.25 | 0.589 |
| *Rassf3* | Spleen | https://genenetwork.org/show_trait?trait_id=10372844&dataset=UTHSC_SPL_RMA_1210F | 10372844 | 7.9 | Chr10: 121.410350 | 0.559 | **0.016** | 0.278 | 0.264 | -0.32 | 0.195 | -0.206 | 0.413 | 0.231 | 0.390 | 0.009 | 0.974 | 0.108 | 0.691 | 0.065 | 0.812 |
| *Rassf3* | White Adipose Tissue | https://genenetwork.org/show_trait?trait_id=TC1000003047.mm.1&dataset=EL_BXDCDScWAT_0216 | TC1000003047.mm.1 | 3.8 | Chr10: 121.973294 | 0.426 | 0.100 | 0.099 | 0.716 | -0.526 | **0.037** | -0.48 | 0.060 | -0.16 | 0.585 | -0.227 | 0.436 | -0.315 | 0.273 | -0.326 | 0.256 |
| *Rassf3* | T-helper Cells | https://genenetwork.org/show_trait?trait_id=1448547_at&dataset=RTHC_0211_R | 1448547_at | 3.9 | Chr10: 121.411254 | 0.793 | 0.060 | 0.771 | 0.072 | -0.662 | 0.152 | -0.429 | 0.397 | 0.735 | 0.157 | 0.6 | 0.285 | -0.13 | 0.834 | 0 | 1.000 |
| *Rassf3* | T-regulatory Cells | https://genenetwork.org/show_trait?trait_id=1448547_at&dataset=RTC_1106_R | 1448547_at | 4.3 | Chr10: 121.411254 | 0.782 | 0.066 | 0.551 | 0.257 | -0.703 | 0.119 | -0.783 | 0.066 | -0.083 | 0.894 | 0.205 | 0.741 | -0.042 | 0.947 | -0.616 | 0.269 |
| *Gns* | Leucocytes | https://genenetwork.org/show_trait?trait_id=ILM3120458&dataset=Illum_BXD_PBL_1108 | ILM3120458 | 1.9 | Chr10: 121.396995 | -0.344 | 0.656 | -0.8 | 0.200 | 0.191 | 0.809 | 0.6 | 0.400 | -0.975 | 0.141 | -1 | **9.00E-11** | 0.64 | 0.558 | 0.5 | 0.667 |
| *Gns* | Neutrophils | https://genenetwork.org/show_trait?trait_id=17237715&dataset=UTHSC-Neut-1014 | 17237715 | 3.1 | Chr10: 121.365090 | 0.331 | 0.423 | 0.167 | 0.693 | -0.452 | 0.261 | -0.286 | 0.493 | -0.036 | 0.939 | -0.036 | 0.939 | -0.314 | 0.493 | -0.107 | 0.819 |
| *Gns* | Spleen | https://genenetwork.org/show_trait?trait_id=10366667&dataset=UTHSC_SPL_RMA_1210 | 10366667 | 4.0 | Chr10: 121.365090 | -0.359 | 0.144 | -0.341 | 0.166 | 0.183 | 0.467 | 0.191 | 0.447 | -0.546 | **0.029** | -0.244 | 0.362 | 0.01 | 0.971 | 0.062 | 0.820 |
| *Gns* | White Adipose Tissue | https://genenetwork.org/show_trait?trait_id=17237715&dataset=EPFLADGL1013 | 17237715 | 4.8 | Chr10: 121.365090 | 0.482 | 0.058 | 0.431 | 0.095 | -0.499 | **0.049** | -0.664 | **0.005** | 0.184 | 0.529 | 0.191 | 0.513 | -0.263 | 0.363 | -0.512 | 0.061 |
| *Gns* | T-helper Cells | https://genenetwork.org/show_trait?trait_id=1433488_x_at&dataset=RTHC_0211_R | 1433488_x_at | 3.4 | Chr10: 121.397110 | -0.588 | 0.220 | -1 | **1.50E-40** | 0.51 | 0.302 | 0.657 | 0.156 | -0.96 | **0.010** | -1 | **1.20E-30** | 0.272 | 0.658 | 0.4 | 0.505 |
| *Gns* | T-regulatory Cells | https://genenetwork.org/show_trait?trait_id=1433488_x_at&dataset=RTC_1106_R | 1433488_x_at | 1.7 | Chr10: 121.397110 | 0.029 | 0.957 | 0.086 | 0.872 | -0.284 | 0.585 | -0.314 | 0.544 | 0.148 | 0.813 | 0.3 | 0.624 | -0.551 | 0.336 | -0.5 | 0.391 |
| *Tbc1d30* | Leucocytes | N/a | N/a | N/a | N/a | N/a | N/a | N/a | N/a | N/a | N/a | N/a | N/a | N/a | N/a | N/a | N/a | N/a | N/a | N/a | N/a |
| *Tbc1d30* | Neutrophils | https://genenetwork.org/show_trait?trait_id=17245466&dataset=UTHSC-Neut-1014 | 17245466 | 3.1 | Chr10: 121.263819 | 0.331 | 0.423 | 0.167 | 0.693 | -0.452 | 0.261 | -0.286 | 0.493 | -0.036 | 0.939 | -0.036 | 0.939 | -0.314 | 0.493 | -0.107 | 0.819 |
| *Tbc1d30* | Spleen | https://genenetwork.org/show_trait?trait_id=10372831&dataset=UTHSC_SPL_RMA_1210 | 10372831 | 10.5 | Chr10: 121.264357 | 0.474 | **0.047** | 0.359 | 0.143 | -0.431 | 0.074 | -0.447 | 0.063 | 0.364 | 0.166 | 0.097 | 0.721 | -0.216 | 0.422 | -0.25 | 0.350 |
| *Tbc1d30* | White Adipose Tissue | https://genenetwork.org/show_trait?trait_id=17245466&dataset=EPFLADGL1013 | 17245466 | 1.9 | Chr10: 121.263819 | 0.386 | 0.140 | 0.069 | 0.799 | -0.272 | 0.308 | -0.025 | 0.927 | -0.151 | 0.605 | -0.253 | 0.383 | 0.057 | 0.848 | 0.275 | 0.342 |
| *Tbc1d30* | T-helper Cells | https://genenetwork.org/show_trait?trait_id=1430607_at&dataset=RTHC_0211_R | 1430607_at | 3.9 | Chr10: 121.266431 | 0.506 | 0.306 | 0.486 | 0.329 | -0.383 | 0.454 | -0.6 | 0.208 | -0.21 | 0.734 | 0.1 | 0.873 | 0.199 | 0.199 | -0.3 | 0.624 |
| *Tbc1d30* | T-regulatory Cells | https://genenetwork.org/show_trait?trait_id=1430607_at&dataset=RTC_1106_R | 1430607_at | 7.9 | Chr10: 121.266431 | 0.665 | 0.150 | 0.886 | **0.019** | -0.719 | 0.107 | -0.771 | 0.072 | 0.788 | 0.113 | 0.8 | 0.104 | -0.558 | **-0.558** | -0.6 | 0.285 |
| *wif1* | Leucocytes | https://genenetwork.org/show_trait?trait_id=ILM7100184&dataset=Illum_BXD_PBL_1108 | ILM7100184 | 2.1 | Chr10: 121.100204 | 0.81 | 0.190 | 0.316 | 0.684 | -0.866 | 0.134 | -0.316 | 0.684 | -0.963 | 0.173 | -0.866 | 0.333 | 0.928 | 0.243 | 0.866 | 0.333 |
| *wif1* | Neutrophils | https://genenetwork.org/show_trait?trait_id=17237701&dataset=UTHSC-Neut-1014 | 17237701 | 2.0 | Chr10: 121.033960 | 0.218 | 0.605 | 0.19 | 0.651 | -0.003 | 0.995 | 0.024 | 0.955 | 0.016 | 0.972 | 0.071 | 0.879 | 0.238 | 0.607 | 0.214 | 0.645 |
| *wif1* | Spleen | https://genenetwork.org/show_trait?trait_id=10366653&dataset=UTHSC_SPL_RMA_1210 | 10366653 | 2.2 | Chr10: 121.034004 | 0.025 | 0.922 | 0.272 | 0.275 | -0.072 | 0.776 | -0.149 | 0.556 | 0.325 | 0.219 | 0.403 | 0.121 | -0.161 | 0.552 | -0.237 | 0.377 |
| *wif1* | White Adipose Tissue | https://genenetwork.org/show_trait?trait_id=17237701&dataset=EPFLADGL1013 | 17237701 | 3.7 | Chr10: 121.033960 | 0.031 | 0.909 | -0.127 | 0.640 | -0.235 | 0.382 | -0.309 | 0.244 | -0.222 | 0.446 | -0.204 | 0.483 | -0.298 | 0.300 | -0.389 | 0.169 |
| *wif1* | T-helper Cells | https://genenetwork.org/show_trait?trait_id=1425425_a_at&dataset=RTHC_0211_R | 1425425_a_at | 4.2 | Chr 10 121.100105 | 0.77 | 0.073 | 0.657 | 0.156 | -0.436 | 0.388 | -0.257 | 0.623 | 0.537 | 0.351 | 0.4 | 0.505 | 0.509 | 0.381 | 0.3 | 0.624 |
| *wif1* | T-regulatory Cells | https://genenetwork.org/show_trait?trait_id=1425425_a_at&dataset=RTC_1106_R | 1425425_a_at | 2.2 | Chr10: 121.100105 | 0.052 | 0.922 | 0.086 | 0.872 | 0.351 | 0.495 | 0.2 | 0.704 | -0.375 | 0.534 | -0.2 | 0.747 | 0.893 | **0.041** | 0.6 | 0.285 |
| *Lemd3* | Leucocytes | N/a | N/a | N/a | N/a | N/a | N/a | N/a | N/a | N/a | N/a | N/a | N/a | N/a | N/a | N/a | N/a | N/a | N/a | N/a | N/a |
| *Lemd3* | Neutrophils | https://genenetwork.org/show_trait?trait_id=17245448&dataset=UTHSC-Neut-1014 | 17245448 | 2.3 | Chr10: 120.923411 | -0.393 | 0.335 | -0.048 | 0.911 | 0.278 | 0.505 | 0.095 | 0.823 | 0.225 | 0.628 | 0.25 | 0.589 | -0.069 | 0.884 | -0.143 | 0.760 |
| *Lemd3* | Spleen | https://genenetwork.org/show_trait?trait_id=10372815&dataset=UTHSC_SPL_RMA_1210 | 10372815 | 2.3 | Chr 10 120.925403 | -0.072 | 0.776 | -0.482 | **0.043** | 0.345 | 0.160 | 0.541 | **0.020** | -0.636 | **0.008** | -0.582 | **0.018** | 0.587 | **0.017** | 0.662 | **0.005** |
| *Lemd3* | White Adipose Tissue | https://genenetwork.org/show_trait?trait_id=17245448&dataset=EPFLADGL1013 | 17245448 | 2.4 | Chr10: 120.923411 | 0.342 | 0.195 | 0.233 | 0.386 | -0.195 | 0.470 | -0.238 | 0.374 | 0.248 | 0.393 | 0.055 | 0.852 | 0.054 | 0.855 | -0.024 | 0.935 |
| *Lemd3* | T-helper Cells | https://genenetwork.org/show_trait?trait_id=1435291_at&dataset=RTHC_0211_R | 1435291_at | 3.9 | Chr10: 120.923477 | -0.734 | 0.097 | -0.714 | 0.111 | 0.899 | **0.015** | 0.829 | **0.042** | -0.586 | 0.299 | -0.5 | 0.391 | 0.852 | 0.067 | 0.7 | 0.188 |
| *Lemd3* | T-regulatory Cells | https://genenetwork.org/show_trait?trait_id=1435291_at&dataset=RTC_1106_R | 1435291_at | 4.6 | Chr10: 120.923477 | 0.621 | 0.189 | 0.6 | 0.208 | -0.657 | 0.156 | -0.943 | **0.005** | 0.326 | 0.593 | 0.3 | 0.624 | -0.373 | 0.536 | -0.9 | **0.037** |
| *Msrb3* | Leucocytes | https://genenetwork.org/show_trait?trait_id=ILM510014&dataset=Illum_BXD_PBL_1108 | ILM510014 | 3.7 | Chr10: 120.791445 | 0.513 | 0.487 | 0.4 | 0.600 | -0.524 | 0.476 | -0.2 | 0.800 | -0.604 | 0.587 | -0.5 | 0.667 | 0.964 | 0.171 | 1 | **9.00E-11** |
| *Msrb3* | Neutrophils | https://genenetwork.org/show_trait?trait_id=17245433&dataset=UTHSC-Neut-1014 | 17245433 | 2.4 | Chr10: 120.781096 | 0.232 | 0.580 | 0.405 | 0.320 | -0.326 | 0.430 | -0.357 | 0.385 | 0.275 | 0.551 | 0.25 | 0.589 | -0.283 | 0.538 | -0.179 | 0.702 |
| *Msrb3* | Spleen | https://genenetwork.org/show_trait?trait_id=10372807&dataset=UTHSC_SPL_RMA_1210 | 10372807 | 2.2 | Chr10: 120.781100 | -0.211 | 0.400 | -0.223 | 0.374 | 0.132 | 0.601 | 0.091 | 0.720 | -0.207 | 0.441 | -0.182 | 0.499 | 0.012 | 0.966 | 0.018 | 0.948 |
| *Msrb3* | White Adipose Tissue | https://genenetwork.org/show_trait?trait_id=17245433&dataset=EPFLADGL1013 | 17245433 | 2.8 | Chr10: 120.781096 | 0.165 | 0.541 | 0.29 | 0.276 | -0.228 | 0.396 | -0.409 | 0.115 | 0.332 | 0.246 | 0.328 | 0.253 | -0.215 | 0.459 | -0.427 | 0.128 |
| *Msrb3* | T-helper Cells | https://genenetwork.org/show_trait?trait_id=1439151_at&dataset=RTHC_0211_R | 1439151_at | 2.7 | Chr10: 120.783223 | -0.726 | 0.102 | -0.543 | 0.266 | 0.505 | 0.307 | 0.086 | 0.872 | -0.283 | 0.644 | -0.2 | 0.747 | -0.271 | 0.659 | -0.6 | 0.285 |
| *Msrb3* | T-regulatory Cells | https://genenetwork.org/show_trait?trait_id=1439151_at&dataset=RTC_1106_R | 1439151_at | 2.3 | Chr10: 120.783223 | 0.729 | 0.100 | 0.829 | **0.042** | -0.537 | 0.272 | -0.486 | 0.329 | 0.868 | 0.056 | 0.7 | 0.188 | -0.002 | 0.997 | -0.1 | 0.873 |
| *Cand1* | Leucocytes | N/a | N/a | N/a | N/a | N/a | N/a | N/a | N/a | N/a | N/a | N/a | N/a | N/a | N/a | N/a | N/a | N/a | N/a | N/a | N/a |
| *Cand1* | Neutrophils | https://genenetwork.org/show_trait?trait_id=17245342&dataset=UTHSC-Neut-1014 | 17245342 | 2.2 | Chr10: 119.198812 | -0.146 | 0.730 | -0.571 | 0.139 | 0.136 | 0.748 | 0.381 | 0.352 | -0.299 | 0.515 | -0.571 | 0.180 | 0.098 | 0.835 | 0.321 | 0.482 |
| *Cand1* | Spleen | https://genenetwork.org/show_trait?trait_id=10372750&dataset=UTHSC_SPL_RMA_1210 | 10372750 | 2.0 | Chr10: 119.202646 | -0.116 | 0.648 | -0.377 | 0.123 | 0.216 | 0.389 | 0.248 | 0.321 | -0.363 | 0.166 | -0.365 | 0.165 | 0.28 | 0.293 | 0.212 | 0.431 |
| *Cand1* | White Adipose Tissue | https://genenetwork.org/show_trait?trait_id=17245342&dataset=EPFLADGL1013 | 17245342 | 1.7 | Chr10: 119.198812 | 0.196 | 0.468 | 0.173 | 0.522 | -0.038 | 0.890 | 0.011 | 0.968 | 0.323 | 0.260 | 0.156 | 0.594 | 0.109 | 0.710 | 0.09 | 0.090 |
| *Cand1* | T-helper Cells | https://genenetwork.org/show_trait?trait_id=1428649_at&dataset=RTHC_0211_R | 1428649_at | 2.5 | Chr10: 119.201927 | 0.87 | **0.024** | 0.6 | 0.208 | -0.797 | 0.058 | -0.943 | **0.005** | 0.158 | 0.800 | 0.3 | 0.300 | -0.194 | 0.754 | -0.9 | **-0.900** |
| *Cand1* | T-regulatory Cells | https://genenetwork.org/show_trait?trait_id=1428649_at&dataset=RTC_1106_R | 1428649_at | 1.9 | Chr10: 119.201927 | 0.356 | 0.489 | -0.143 | 0.787 | -0.318 | 0.538 | -0.2 | 0.704 | -0.931 | **0.021** | -1 | **1.20E-30** | 0.253 | 0.681 | 0.4 | 0.505 |
