## Supplementary Table 6 for "Determining susceptibility loci in triple negative breast cancer using a novel pre-clinical model"

**Supplemental Table 6. Chromosome 4 QTL Protein Coding Genes for Necrosis.** An interval generated in GEMMA for chr4 (103.640707-105.243637 Mb) contains 23 annotated genes and open reading frames (ORFs), of which 6 were protein coding with gene symbol, description, start point, length, and variant or single nucleotide polymorphims (SNP) counts and density reported.

| Symbol | Gene Description | Mb Start | Length (Kb) | SNP Count | SNP Density |
|---|---|---|---|---|---|
| *Dab1* | disabled 1 | 103.619500 | 1125.34 | 1398 | 1.24 |
| *C8b* | complement component 8, beta polypeptide | 104.766317 | 38.23 | 0 | 0.00 |
| *C8a* | complement component 8, alpha polypeptide | 104.815679 | 60.81 | 2 | 0.03 |
| *Fyb2* | FYN binding protein 2 | 104.835301 | 181.56 | 5 | 0.03 |
| *Prkaa2* | protein kinase, AMP-activated, alpha 2 catalytic subunit | 105.029649 | 80.25 | 141 | 1.76 |
| *Plpp3* | phospholipid phosphatase 3 | 105.157347 | 75.42 | 49 | 0.65 |
