## Supplementary Table 7 for "Determining susceptibility loci in triple negative breast cancer using a novel pre-clinical model"

**Supplemental Table 7. Chromosome 12 QTL Protein Coding Genes for Stroma and Vascularity.** An interval generated in GEMMA for chr12 64.387817- 69.575599 Mb) contains 172 annotated genes and open reading frames (ORFs), of which 43 were protein coding with gene symbol, description, start point, length, and variant or single nucleotide polymorphims (SNP) counts and density reported.

| Symbol | Gene Description | Mb Start | Length (Kb) | SNP Count | SNP Density |
|---|---|---|---|---|---|
| *Mia2* | MIA SH3 domain ER export factor 2 | 59.095582 | 94.64 | 22 | 0.23 |
| *Fbxo33* | F-box protein 33 | 59.200655 | 21.31 | 11 | 0.52 |
| *Lrfn5* | leucine rich repeat and fibronectin type III domain containing 5 | 61.522851 | 335.08 | 70 | 0.21 |
| *Spanxn4* | SPANX family, member N4 | 62.687787 | 0.55 | 0 | 0.00 |
| *Fscb* | fibrous sheath CABYR binding protein | 64.471333 | 3.57 | 0 | 0.00 |
| *Klhl28* | kelch-like 28 | 64.941486 | 24.10 | 0 | 0.00 |
| *Togaram1* | TOG array regulator of axonemal microtubules 1 | 64.965576 | 57.00 | 3 | 0.05 |
| *Fkbp3* | FK506 binding protein 3 | 65.062432 | 11.51 | 0 | 0.00 |
| *Fancm* | Fanconi anemia, complementation group M | 65.074110 | 57.73 | 2 | 0.03 |
| *Mis18bp1* | MIS18 binding protein 1 | 65.132734 | 39.85 | 0 | 0.00 |
| *Wdr20rt* | WD repeat domain 20, retrogene | 65.225517 | 2.94 | 0 | 0.00 |
| *Rpl10l* | ribosomal protein L10-like | 66.283379 | 1.02 | 0 | 0.00 |
| *Mdga2* | MAM domain containing glycosylphosphatidylinositol anchor 2 | 66.459419 | 763.13 | 91 | 0.12 |
| *Rps29* | ribosomal protein S29 | 69.157722 | 1.46 | 0 | 0.00 |
| *Lrr1* | leucine rich repeat protein 1 | 69.168814 | 10.20 | 7 | 0.69 |
| *Rpl36al* | ribosomal protein L36A-like | 69.182734 | 1.33 | 0 | 0.00 |
| *Mgat2* | mannoside acetylglucosaminyltransferase 2 | 69.184158 | 2.62 | 0 | 0.00 |
| *Dnaaf2* | dynein, axonemal assembly factor 2 | 69.189087 | 11.00 | 1 | 0.09 |
| *Pole2* | polymerase (DNA directed), epsilon 2 (p59 subunit) | 69.201773 | 26.43 | 8 | 0.30 |
| *Klhdc1* | kelch domain containing 1 | 69.227729 | 56.92 | 17 | 0.30 |
| *Klhdc2* | kelch domain containing 2 | 69.296681 | 14.01 | 4 | 0.29 |
| *Nemf* | nuclear export mediator factor | 69.311541 | 45.66 | 1 | 0.02 |
| *Arf6* | ADP-ribosylation factor 6 | 69.372150 | 3.83 | 0 | 0.00 |
| *Vcpkmt* | valosin containing protein lysine (K) methyltransferase | 69.576735 | 6.28 | 12 | 1.91 |
| *Sos2* | SOS Ras/Rho guanine nucleotide exchange factor 2 | 69.583761 | 98.09 | 219 | 2.23 |
| *L2hgdh* | L-2-hydroxyglutarate dehydrogenase | 69.690436 | 34.44 | 76 | 2.21 |
| *Dmac2l* | distal membrane arm assembly complex 2 like | 69.724913 | 19.75 | 24 | 1.22 |
| *Cdkl1* | cyclin-dependent kinase-like 1 (CDC2-related kinase) | 69.745167 | 46.03 | 63 | 1.37 |
| *Map4k5* | mitogen-activated protein kinase kinase kinase kinase 5 | 69.803757 | 89.41 | 238 | 2.66 |
| *Atl1* | atlastin GTPase 1 | 69.893105 | 70.98 | 181 | 2.55 |
| *Sav1* | salvador family WW domain containing 1 | 69.965009 | 23.13 | 56 | 2.42 |
| *Nin* | ninein | 70.011435 | 102.38 | 403 | 3.94 |
| *Abhd12b* | abhydrolase domain containing 12B | 70.154124 | 35.97 | 128 | 3.56 |
| *Pygl* | liver glycogen phosphorylase | 70.190815 | 36.87 | 141 | 3.82 |
| *Trim9* | tripartite motif-containing 9 | 70.244533 | 103.34 | 104 | 1.01 |
| *Tmx1* | thioredoxin-related transmembrane protein 1 | 70.453154 | 14.47 | 3 | 0.21 |
| *Frmd6* | FERM domain containing 6 | 70.825456 | 76.78 | 10 | 0.13 |
| *Actr10* | ARP10 actin-related protein 10 | 70.937857 | 26.86 | 0 | 0.00 |
| *Psma3* | proteasome subunit alpha 3 | 70.969283 | 26.59 | 2 | 0.08 |
| *Arid4a* | AT rich interactive domain 4A (RBP1-like) | 71.015081 | 84.27 | 11 | 0.13 |
| *Tomm20l* | translocase of outer mitochondrial membrane 20-like | 71.111428 | 11.79 | 5 | 0.42 |
| *Timm9* | translocase of inner mitochondrial membrane 9 | 71.123172 | 13.54 | 26 | 1.92 |
| *Dact1* | dishevelled-binding antagonist of beta-catenin 1 | 71.309884 | 10.22 | 48 | 4.70 |
